# Quantifying nanotherapeutics penetration using hydrogel based microsystem as a new 3D *in vitro* platform

**DOI:** 10.1101/2021.01.17.427020

**Authors:** Saba Goodarzi, Audrey Prunet, Fabien Rossetti, Guillaume Bort, Olivier Tillement, Erika Porcel, Sandrine Lacombe, Ting-Di Wu, Jean-Luc Guerquin-Kern, Hélène Delanoë-Ayari, François Lux, Charlotte Rivière

## Abstract

The huge gap between 2D *in vitro* assays used for drug screening, and the *in vivo* 3D-physiological environment hampered reliable predictions for the route and accumulation of nanotherapeutics *in vivo.* For such nanotherapeutics, Multi-Cellular Tumour Spheroids (MCTS) is emerging as a good alternative *in vitro* model. However, the classical approaches to produce MCTS suffer from low yield, slow process, difficulties in MCTS manipulation and compatibility with high-magnification fluorescent optical microscopy. On the other hand, spheroid-on-chip set-ups developed so far require a microfluidic practical knowledge difficult to transfer to a cell biology laboratory.

We present here a simple yet highly flexible 3D-model microsystem consisting of agarose-based microwells. Fully compatible with the multi-well plates format conventionally used in cell biology, our simple process enables the formation of hundreds of reproducible spheroids in a single pipetting. Immunostaining and fluorescent imaging including live high-resolution optical microscopy can be performed *in-situ*, with no manipulation of spheroids.

As a proof-of-principle of the relevance of such *in vitro* platform for nanotherapeutics evaluation, this study investigates the kinetic and localization of nanoparticles within colorectal cancer MCTS cells (HCT-116). The nanoparticles chosen are sub-5 nm ultrasmall nanoparticles made of polysiloxane and gadolinium chelates that can be visualized in MRI (AGuIX^®^, currently implicated in clinical trials as effective radiosensitizers for radiotherapy) and confocal microscopy after addition of Cy 5.5. We show that the amount of AGuIX^®^ nanoparticles within cells is largely different in 2D and 3D. Using our flexible agarose-based microsystems, we are able to resolve spatially and temporally the penetration and distribution of AGuIX^®^ nanoparticles within MCTS. The nanoparticles are first found in both extracellular and intracellular space of MCTS. While the extracellular part is washed away after few days, we evidenced intracellular localisation of AGuIX^®^, mainly within lysosomes compartment, but also occasionally within mitochondria. Our agarose-based microsystem appears hence as a promising 3D *in vitro* user-friendly platform for investigation of nanotherapeutics transport, ahead of *in vivo* studies.

Graphical abstract

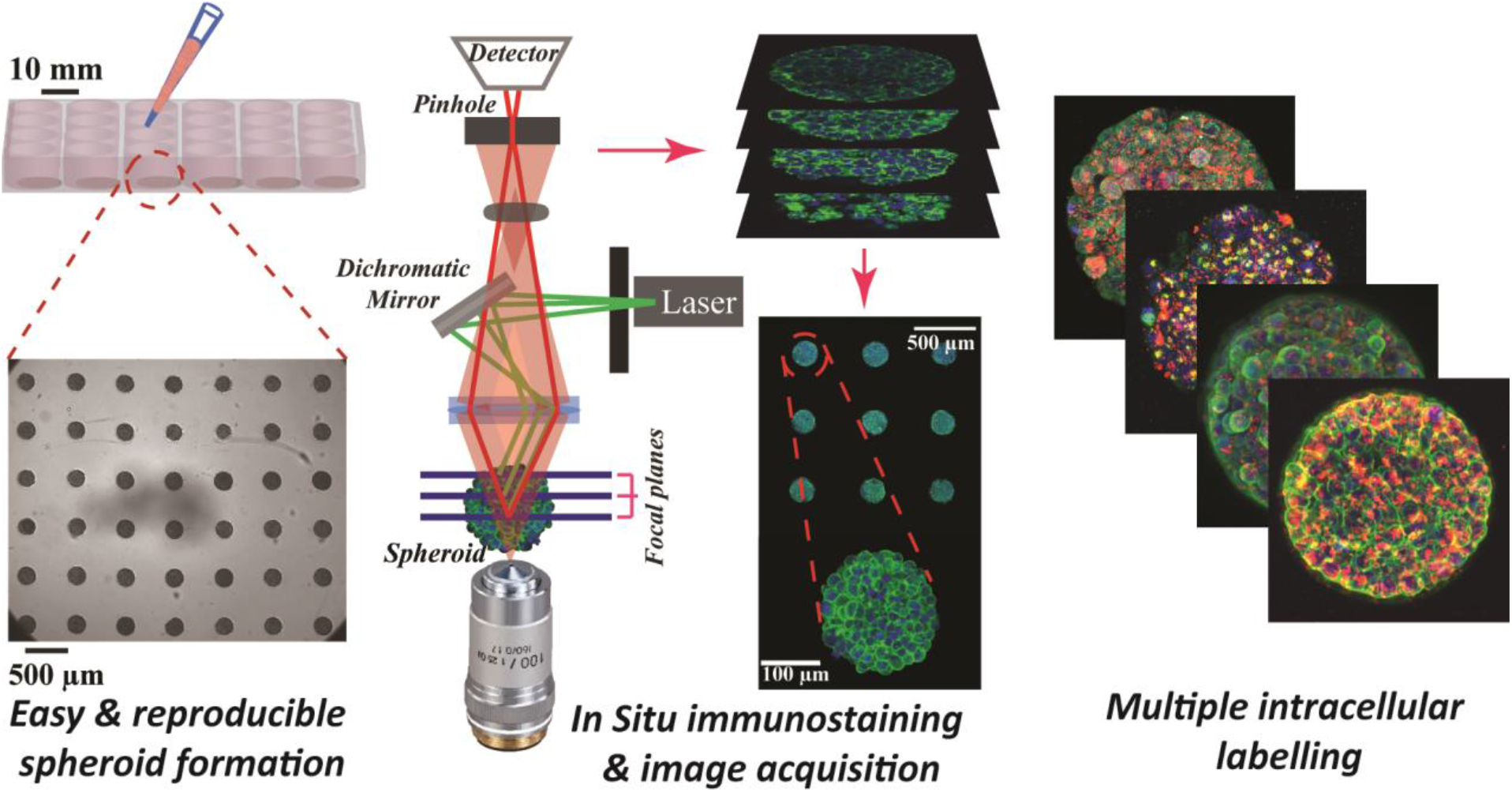

## Introduction

There is an ongoing effort to develop efficient therapeutics for cancer treatment including nano-drugs and nanoparticles, nevertheless, the clinical translation of these therapeutics has to overcome numerous challenges from early stages of development to a successful translation ^1,2^. Currently, the standard pipeline for drug development is the following: (1) efficacy tests on 2D *in vitro* assays, and (2) on rodent *in vivo* models, (3) regulatory toxicity tests on two animal species and (4) clinical trials.

However, 2D *in vitro* assays do not replicate the 3D-physiological environment encountered by the cells *in vivo*. That could be the underlying reason of the high rate of clinical failure in development of new drugs. On the other hand, there is also a rising questioning of the economical and ethical relevance of rodent animal models, in particular because such models are not fully representative of human specificity ^3^. Getting as close as possible to the *in vivo* situation in *in vitro* models is a key issue to truly understand and control cancer cell response, accompanied by reduction in animal usage. For the pharmaceutical industry, tackling this issue will enable better identification of relevant therapeutics by performing relevant screening on 3D models. For precision medicine, it will help physicians to adjust the therapeutic treatment, in complement of current clinical analysis ^2,4^. For fundamental research, it will allow deciphering cell response in a truly relevant context.

Many approaches have been developed during the past decade to set-up various organ-on-a-chip or tumour-on-a-chip devices, integrating many different *in vivo* features in a miniaturized *in vitro* format ^5,6^. This is particularly important for emerging nanosized therapeutics ^7^. The presence of different physiological barriers such as cell-cell compaction, tumour heterogeneity, dense extracellular matrix along with various cancer-associated cells, will decrease the amount of nanotherapeutics effectively reaching the targeted tumour cells ^8,9^. The lack of such physiological context hampered reliable predictions for the route and accumulation of those nanoparticles *in vivo* ^10^ and is a major limitation for the efficient development of novel therapeutic approaches ^11^. To move beyond the classical 2D-plastic dishes, different 3D *in vitro* models have been developed to try to better replicate *in vivo* complexity of tumour microenvironment ^12^. Among them, Multi-Cellular Tumour Spheroids (MCTS) recapitulate many tumour features including 3D cellular architecture, cellular heterogeneity, signalling pathways and physiochemical gradient similar to real *in vivo* tumour micrometastasis (for spheroids > 500 μm in diameter) ^13–17^.

MCTS could be prepared with various techniques ^13^ such as using non-adherent surfaces ^18^, spinner flasks ^19^ or hanging drop methods ^20^. Emerging attempts to integrate spheroids in microfluidic set-ups open up new possibilities to deal with the low yield and slow process of the classical approach ^21,22^. However, such spheroid-on-chip approaches require a microfluidic practical knowledge that is difficult to transfer to a cell biology laboratory.

In addition, the polymeric materials commonly used for such devices (Polydimethylsiloxane -PDMS) suffers from major limitations, precluding its usage for efficient drug screening in physiological conditions ^23^: large absorption of therapeutics ^24,25^ (resulting in the underestimation of cell response to drugs), non-permeability to small water-soluble molecules (leading to fast-medium conditioning if continuous flow is not provided otherwise), rigidity several orders of magnitude larger than physiological condition (MPa *vs* kPa range *in vivo* ^26^).

To go beyond PDMS and its limitations, hydrogel-based microwells devices have been considered ^27,28^. Hydrogels are network of cross-linked polymers with tuneable physical properties and high capacity of water retaining and interconnected pores enabling free diffusion of O_2_, nutrient and metabolic wastes, which make them favourable alternatives in micro-system applications. Various techniques using natural or synthetic hydrogels for MCTS formation, have been developed ^28–33^. However, none of these set-ups meets all the criteria required for long-term time-lapse analysis (i.e. compatibility with high-resolution video-microscopy, efficient medium and oxygen renewal, *in-situ* immunostaining/drug application, no reduction of the available drug dose, easy cell retrieval for further standard molecular analyses), within a physiological stiffness range.

We present here a simple yet highly flexible 3D-model microsystem consisting of agarose-based microwells. This hydrogel with tuneable rigidity and great integrity presents several advantages making it a suitable biomaterial in cell studies ^34,35^. The tuneable mechanical properties of the agarose can reproduce the *in vivo* microenvironment stiffness. Its porous nature enables the free diffusion of salt and small chemical species (hydrodynamic diameter <30 nm in 2% agarose ^36^, which is the case for most proteins). Our simple process enables the formation of hundreds of reproducible spheroids in a single pipetting, and its compatibility with multi-well plate formats conventionally used in cell biology can accelerate the screening of drugs in comparison with conventional 3D models. Of note, these microwells can also be manufactured on coverslips, opening the possibility for live high-resolution optical microscopy. In addition, the hydrogel-based microwells provides a user-friendly platform for *in-situ* immunostaining and can be used for in-depth analysis of cell phenotypic modifications after drug treatment.

As a proof-of-principle of the relevance of such *in vitro* platform for the evaluation of nanoparticles screening, the aim of this study was to analyse the kinetic and localization of these nanoparticles within colorectal cancer cells MCTS (HCT-116). The nanoparticles chosen for this proof-of-concept study are sub-5 nm ultrasmall nanoparticles made of polysiloxane and gadolinium (Gd) chelates that can be visualized in MRI and confocal microscopy (after functionalization by Cy5.5, a near-infrared fluorophore). These nanoparticles, called AGuIX^®^, are effective radiosensitizers for radiotherapy ^37^ and are now implicated in three clinical trials associating radiotherapy with AGuIX^®^ for treatment of multiple brain metastases by whole brain radiation therapy (NanoRad 2, Phase II, multicentric), stereotactic radiosurgery (NanoStereo, Phase II, multicentric) and cervical cancer (Phase Ib, Gustave Roussy). Nanoparticles-cell interactions and internalization pathways of these nanoparticles have been assessed *in vitro* in 2D ^38^, but never in 3D multicellular tumour spheroids.

We show in this study that the 3D cell arrangement highly impacts the amount of AGuIX^®^ nanoparticles within cells. Using our flexible agarose-based microsystem, we were able to resolve spatially and temporally the penetration and distribution of AGuIX^®^ nanoparticles within tumour spheroids. The nanoparticles were first found in both extracellular and intracellular space of spheroids, mostly within lysosomes compartment, but also occasionally within mitochondria. Whereas the extracellular part was washed away after few days, the colocalization with lysosomes remained almost constant. Our agarose-based microsystem appears hence as a promising 3D *in vitro* platform for investigation of nanotherapeutics transport, ahead of *in vivo* studies.

## Materials and Methods

### Hydrogel based microsystem

Agarose-based microsystems were prepared using moulding procedures. First, a silicon wafer moulds was made using classical photolithography technique (**Fig. 1 A**). The mould consists of an array of 130 cylindrical wells of 200 μm in diameter, and 250 μm in height), created using the SU8-2100 photosensitive resin.

**Figure.1.**
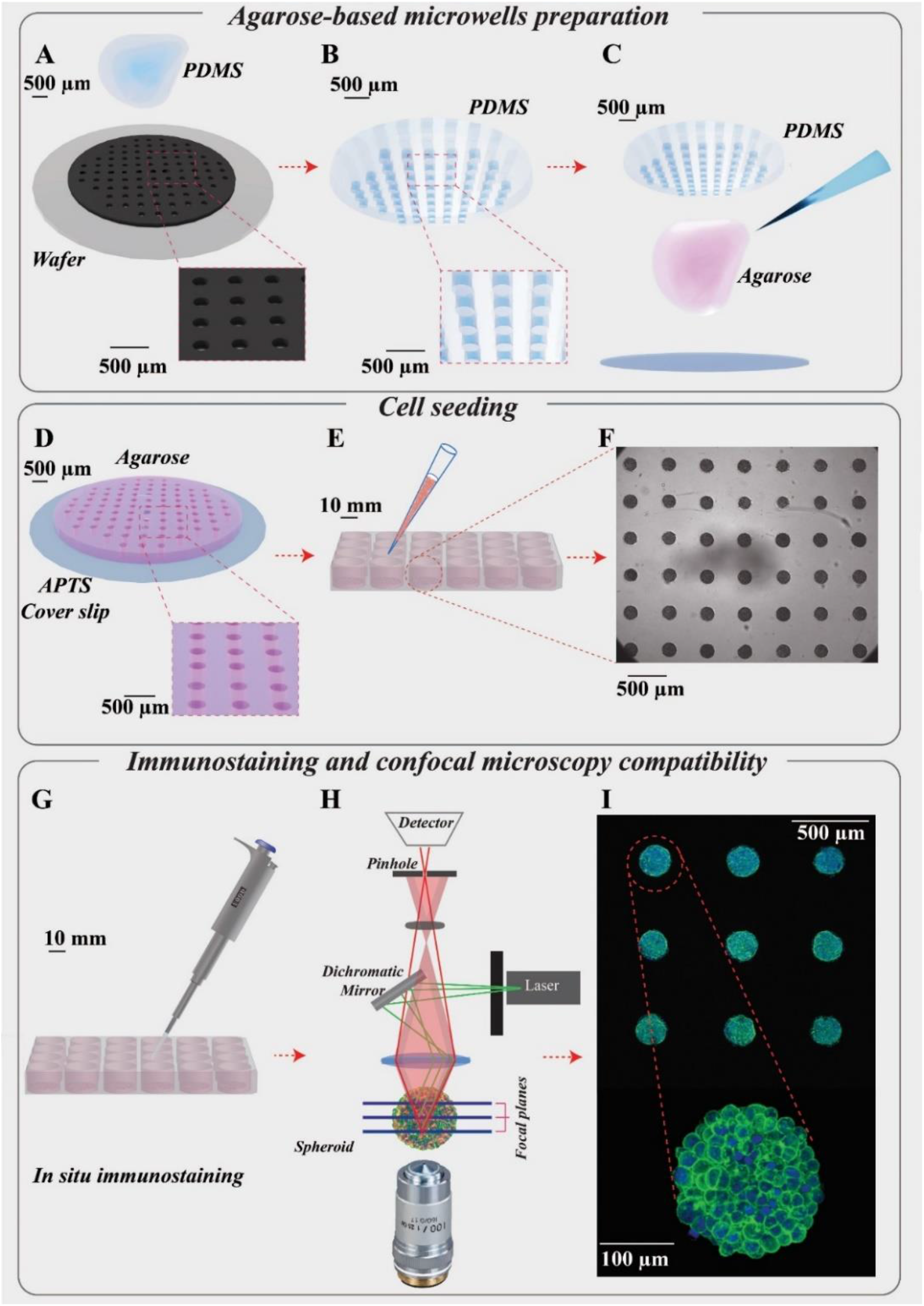
Presentation of the hydrogel-based microsystems for spheroids growth and follow-up. **(A)** Silicon wafer mould made by photolithography. **(B)** PDMS replica mould made from the silicon wafer mould. **(C)** Moulding of agarose using PDMS replica moulds on a cover slip functionalized by APTS to make the agarose microsystem adhesive on the cover slip. **(D)** Cylindrical agarose microwells with diameter and height of 200 μm for each microwell. **(E)** Cell seeding using agarose microsystem in a 24-well plate for preparation of spheroids (leading to the formation of 130 spheroids/well). **(F)** Optical microscopy images of several homogenous HCT116 cell spheroids made in agarose microsystem (5X magnification) at day 6. **(G)** In-situ immunostaining of spheroids in microwells in a 24-well plate. **(H)** Diagram of confocal fluorescence microscopy. **(I)** Maximal Image Projection (MIP) of confocal fluorescence images of spheroids in agarose microsystem labelled for actin (green) and nuclei (blue) (10X magnification) and an enlarged MIP of one of the spheroids (20X magnification).

A polydimethylsiloxane (PDMS) replica mould was then casted on this master mould (**Fig. 1 B)** and used for agarose moulding. The agarose moulding procedure was differing depending on the aim of the experiments: (1) for imaging of fixed samples, the microwells were free-standing in each well of a multi-well plate, enabling easy retrieval and transfer (see detailed description below); (2) for time-lapse imaging, agarose moulding is performed on 3-AminoPropylTriethoxiSilane (APTS)-functionalised coverslips, enabling to directly bond the microwells to the coverslips and avoiding any drift during acquisition (**Fig. 1 D,** patented process^39^)

Agarose solution (2%, w/v) was prepared by dissolving ultra-pure agarose powder (Invitrogen™) in water. Autoclave was used for the dissolution to avoid bubbles formation (121°C, 15 min).

#### Moulding of free-standing microwells

The agarose solution (300μL) was deposited on a warmed PDMS mould (at 78°C) and a coverslip was then placed on top of the drop of agarose to spread it with a constant thickness on the mould. After agarose gelation into the desired shape (10 min), the coverslip was removed and the moulded agarose microwells were cut to fit in the wells of a 24 multi-well plate. The microwells were then placed in a 24-multi well plate and kept hydrated with PBS (1 mL/well). The plate was UV-sterilized (8 W, 254 nm) for 20 min on opened and closed state and kept at 4°C until used. The day before each experiment, PBS was replaced by culture medium and let to diffuse within each microwell by overnight incubation at 37°C before cell seeding.

#### Moulding on APTS-functionalised coverslips

First, holes were drilled in each well of a 12-well plate (diameter 16 mm) to prepare the plate for the coverslips. Round coverslips (diameter 20 mm) were incubated in a 1% APTS-5mM acetic acid solution (Acros ref 43094100 for APTS, vwr ref 20104298 for acetic acid) for 20 min under stirring condition. Coverslips were then extensively rinsed with water and dried on a hot plate (100 °C, 15 min). Such APTS-functionalised coverslips are then used immediately for agarose moulding using the same procedure as the one described above for free-standing microwells. After agarose gelation, the PDMS mould was removed, the agarose microwells remaining attached to the APTS-functionalised coverslip. These coverslips with microwells were glued to the 12-well plate using curing optical adhesive (Norland products, NOA 81) activated by 30 seconds of exposure to a UV lamp (12 W-365 nm). The plate was then UV-sterilized using the same procedure as the one described above for free-standing microwells.

### Colorectal cancer cell line, HCT116 and culture condition

HCT-116 colorectal carcinoma (CCL-247) cell line was purchased from the American Type Culture Collection (ATCC, Virginia, USA). All cells were cultured in Dulbecco’s Modified Eagle’s medium (DMEM-Glutamax, Gibco™), supplemented with 10% of heat-inactivated Fetal Bovine Serum (FBS; Sigma, St. Louis, Missouri, US), 100 units/100 μg of penicillin/streptomycin (Gibco™).

Routinely, the HCT-116 cells were grown in T-25 cell culture flasks and were placed in the incubator at 37°C with a 5% CO_2_ atmosphere. The culture medium was changed regularly, and the cell passage was carried out at 70% confluency every 3 days. The cell passage was performed using recombinant cell-dissociation enzyme (TrypLE, Gibco™) to detach cells followed by neutralizing with culture medium. The cell suspension was centrifuged at 1000 rpm (equal to 106 g) for 5 min, the supernatant was discarded, and the cell pellet was resuspended in 1 mL. The number of cells was counted with a Neubauer chamber, and final cell volume was adjusted to reach the desired cell concentration.

### Multicellular tumour spheroids

MCTS of HCT-116 cells were formed in 24-well plates containing agarose microwells in each well. After trypsinization and centrifugation, 120,000 cells in 1 mL complete medium was added in each well (containing each 1 microsystem). To encourage and accelerate cell aggregation, the 24-well plate was placed under orbital agitation (160 rpm) for 15 min in the incubator at 37°C and 5% CO_2_. After 4 h, the plate was rinsed with fresh medium to remove cells that did not reach the microwells. After 2 days, spheroids were ready for incubation with nanoparticles.

### Monolayer cell culture

After trypsinization and centrifugation of HCT-116 cells in culture, a cell suspension with 120,000 cells in 1 ml was prepared. The cell suspension was added to tissue-treated coverslip plates (either 300 μL in 8-well Ibidi^®^ or 2mL in 12-well plate). Cells were incubated with nanoparticles 48 h after cell seeding.

### Preparation of Cy5.5 conjugated Gadolinium based nanoparticles (AGuIX^®^-Cy5.5)

The Gd-based nanoparticles (AGuIX^®^) synthesized by NH TherAguix (Lyon, France) are composed of a polysiloxane matrix surrounded by covalently bound DOTAGA-Gd ((1,4,7,10-tetraazacyclodode-cane-1-glutaric acid-4,7,10-triacetic acid)-Gd). The synthesis process is already described in the literature ^40^. Briefly, AGuIX^®^ nanoparticles are composed of a polysiloxane network surrounded by Gd chelates. The chemical composition of AGuIX^®^ nanoparticles is (GdSi_6.5_N_6_C_25_O_21_H_42_, 10 H_2_O)_n_ with a molar mass around 10 kDa. The hydrodynamic diameter of the AGuIX^®^ nanoparticles is close to 5 nm; and the AGuIX^®^ nanoparticles are characterized by a zeta potential of 9.0 ± 5.5 mV at pH 7.2. These AGuIX^®^ nanoparticles were further conjugated to Cyanine-5.5(Cy5.5) fluorophore to make them detectable by confocal fluorescence microscopy. They are referred as AGuIX^®^-Cy5.5 nanoparticles in the rest of the article.

### Incubation of cells with AGuIX^®^-Cy5.5 nanoparticles

To incubate MCTS and monolayer cells with AGuIX^®^-Cy5.5 nanoparticles, an intermediate solution of AGuIX^®^ -Cy5.5 nanoparticles with 100 mM concentration of Gd was prepared in distilled-water. From this intermediate solution, just before the incubation with cells, AGuIX^®^-Cy5.5 solutions were prepared in fresh DMEM with Gd concentrations of 0.8, 1.5 and 2 mM respectively. The MCTS in all microsystems of a 24-well plate were incubated with 1 mL of AGuIX^®^-Cy5.5 nanoparticles solution. For cell monolayers, an Ibidi^®^ 8-well plate or a 12-wells plate was used, and cells were incubated with 200 μL or 2 mL AGuIX^®^-Cy5.5 solution, respectively.

### Inductively coupled plasma-mass spectrometry (ICP-MS)

Concentrations of Gd were analysed using a validated inductively coupled plasma-mass spectrometry (ICP-MS). To prepare samples for this analysis, spheroids and monolayer cultured cells were incubated with AGuIX^®^-Cy5.5 nanoparticles with 0.8, 1.5 and 2 mM concentration in Gd for 24 h. After incubation, spheroids were rinsed three times with PBS for 15 min each and dissociate using Trypsin + EDTA (Gibco). The number of cells in each microwells was evaluated using a Neubaeur chamber. The cell suspensions in trypsin + EDTA of each sample were then centrifuged (900 g for 5 min), the supernatants discarded, and cells pellets were dissolved in 150 μl HNO3 69% (ROTH) at 80° C for 3 h. The volume of samples was adjusted to 10 mL by adding ultra-pure water and Gd concentration in each sample was measured using ICP-Mass Spectrometer (PerkinElmer, NexION^®^ 2000). A similar procedure was used for monolayer cell culture (the cells were rinsed with PBS (3x 5 min) and detached using trypsin (Gibco)).

The mean value of cell volume was calculated by measuring the cell diameter after detaching or dissociation using bright field microscopy followed by image processing using ImageJ software ^41^. Accordingly, Gd concentrations obtained by ICP-MS measurements were divided by the average cell volume calculated.

### Localization of nanoparticles: Fixation, permeabilization and immunostaining

First, cell nuclei and actin filaments in cytoskeleton were labelled. After incubation with AGuIX^®^-Cy5.5 nanoparticles, spheroids were rinsed with PBS (3x 5 min), then fixed in paraformaldehyde (4%) for 20 min and permeabilized using 0.1% Triton X-100 (Acros) for 10 min. After blocking with 3% bovine serum albumin (BSA, Sigma-Aldrich) for 20 min, samples were incubated with phalloidin-546 solution (Invitrogen™, A22283, 1:50 in PBS) containing nucgreen™-Dead 488 (Invitrogen™, R37109, 1 drop/5ml in PBS) at 4°C overnight. The procedure ended with rinsing spheroids with PBS (3x 5 min).

In a second series of experiments, to find out the precise intracellular localization of AGuIX^®^-Cy5.5 nanoparticles, three antibodies were used to label the main cell compartments: EEA1 for early endosomes (CellSignaling Technology, #3288), AIF for mitochondria (CellSignaling Technology, # 5318) and LAMP-1 for lysosomes (CellSignaling Technology, #9091). After fixation in paraformaldehyde (4%) for 20 min and rinsing with PBS (3x 5 min), according to the protocol proposed by the manufacture, cells were blocked in the buffer (PBS/ 5% BSA/ 0.3% Triton™ X-100) for 60 min, and rinsed with PBS (3x 5 min). These samples, either spheroids in microwells, either cell monolayer in Ibidi plates were incubated with EEA1 (1:100), AIF (1:400) and LAMP1 (1:200) in the buffer (PBS/ 1% BSA/ 0.1% Triton™ X-100) overnight.

The incubation buffers were aspirated and cells were rinsed with PBS (3x 5 min). For the secondary antibody, Goat-anti rabbit IgG-Alexa 555 (Invitrogen™, A21428, 1:500, in PBS/ 1% BSA/ 0.1% Triton™ X-100) was used. All samples were then incubated with nucgreen™ Dead 488 (Invitrogen™, 1 drop/5 ml in PBS) overnight for spheroids and 4 h for cell monolayers. In the last step, they were rinsed with PBS (3×5 min).

### Spheroids Clarification

Optical imaging of three-dimensional biological samples can be performed using confocal fluorescence microscopy which images these 3D samples via optical sectioning. However this technique faces several limitations including, light scattering, attenuation of photon due to light absorption and local refractive index differences limiting the light depth of penetration ^42^. Many different clarification techniques have been developed to overcome such issues ^43,44^. In the current study, the clearing efficiency of two methods has been analysed using nucgreen™ signals in HCT-116 spheroids: RapiClear 1.52 (SunjinLab) and Glycerol ^45^. Based on the quantification of fluorescence intensities (**Fig.SI 1**), clarification with glycerol/PBS (80%/20%) has been chosen to clear spheroids in this study.

The solution for clarifying spheroids was prepared by mixing glycerol (99.5%, VWR Chemicals) with PBS by the ratio of (80%/20%). A fresh solution was prepared for every experiment. To clarify spheroids, just after fixation, they were incubated in glycerol solution for 24 h. A detailed description of the mounting procedure used for imaging of live and fixed spheroids is described in Supplementary **Fig.SI 2.** For most experiments, the microsystems were incubated with a fresh glycerol solution and mounted between 2 coverslips, separated by a 1 mm sticky spacer (2×0.5mm thick Ispacer, SunJin Lab).

### Confocal fluorescence microscopy

Image acquisitions of spheroids and cell monolayers were carried out with a confocal microscope (Leica SP5) using either a 20X dry objective (NA= 0.7), a 25X water immersion objective (NA=0.95), or a 40X oil immersion objective (NA=1.25). Image acquisitions in Z direction was performed using a 1 μm z-step. Automatic image acquisitions for a large number of spheroids were performed (about 4 h for 30 spheroids using 30% power for AGuIX^®^-Cy5.5 nanoparticles (λ_excitation_ = 633 nm)).

### Image processing

Images obtained by confocal fluorescence microscopy were analysed using a dedicated Matlab routine. While spheroids were imaged using optical sectioning in Z direction, it was useful to quantify the average signal intensity along the radius of each spheroid.

To do this, the entire surface of each spheroid, at each imaging depth were first segmented using the intensity signals coming from every nuclei (labelled with nucgreen™-488). From this segmentation, the segmented spheroids slices were first fitted into a perfect circle for each imaging depth, followed by fitting each spheroid z-stack into a perfect sphere. By changing the coordinates of analysis from Cartesian (x, y, z) to spherical (R, theta, phi) coordinates, the mean intensity of AGuIX^®^-Cy5.5 nanoparticles was averaged along theta and phi angles. The obtained averaged intensity was normalised with the maximum grey value of images obtained and plotted as a function of the distance from the periphery.

For cell monolayers, the maximum Z-projection of each field of view imaged by confocal microscopy (obtained by image J) was used and analysed with a Matlab script to quantify the mean intensity in these images. For each sample, the average of the mean intensity computed in the different fields of view was calculated.

### Colocalization quantification

To quantify the colocalization of AGuIX^®^-Cy5.5 nanoparticles with cell organelles from confocal fluorescence images, a dedicated routine has been developed to calculate Pearson’s correlation coefficient, indicating the degree of colocalization between fluorophores. Briefly, for each image of the acquired-stack, a mask of the spheroid was automatically defined using the nucleus staining. Correlation between the far-red and red channels (corresponding to AGuIX^®^-Cy5.5 and organelle-immunostaining respectively) was then computed using the corr2 Matlab function. Using this routine, Pearson’s correlation coefficient was calculated in the spheroid area for each image along the Z-direction (same as acquisition).

### NanoSIMS Cellular imaging

To prepare samples for NanoSIMS cellular imaging, HCT-116 cell spheroids were already incubated with 2mM AGuIX^®^ nanoparticles for 72h and then fixed with 2% glutaraldehyde in cacodylate buffer (0.1 M, pH 7.8) for 60 min, followed by rinsing with PBS (3×5min). Samples were then postfixed with 1% osmium tetroxide follow by uranyl acetate staining and gradually dehydrated in ethanol (30% to 100%) and embedded in Epon.

A 0.2 μm relatively thick section was deposited onto clean Si chip and dried in air before being introduced into a NanoSIMS-50 Ion microprobe (CAMECA, Gennevilliers, France) operating in scanning mode ^46,47^. For the present study, a tightly focused Cs^+^ primary ion beam at an impact energy of 16 keV was used to monitor up to five secondary ion species in parallel from the same sputtered volume: ^12^C^−^, ^12^C^14^N^−^, ^28^Si^−^, ^31^P^−^, as well as ^35^Cl^−^. The primary beam steps over the surface of the sample to create images for these selected ion species. The primary beam intensity was 3 pA with a typical probe size of ≈200 nm. The raster size was 60 μm with an image definition of 512×512 pixels. The acquisition was carried out in multiframe mode with a dwell time of 0.5 ms per pixel and 220 frames were recorded. The image processing was performed using the ImageJ software ^41^. Successive image frames were properly aligned using TOMOJ plugin ^48^ with ^12^C^14^N^−^ images as reference to correct the slight image field shift during the 8 h signal accumulation, before a summed image was obtained for each ion species.

## Results and Discussion

A hydrogel-based microsystem was developed to generate uniform-sized multicellular tumour spheroids (**Fig. 1**). The design of these microwells was meant to meet the following goals: 1) making homogenous and uniform cell spheroids 2) increase the throughput in drug screening 3) be compatible with *in-situ* treatment, immunostaining and image acquisition as well as *ex-situ* characterization techniques. First, a silicon wafer mould was designed and made using classical photolithography technique (**Fig. 1A**). From this silicon wafer mould, counter moulds in Polydimethylsiloxane (PDMS) were prepared (**Fig. 1B**), which could be used several times to replicate microwells with agarose hydrogel. To prepare agarose microwells, 2% ultra-pure agarose solution was poured on PDMS moulds (**Fig. 1C**) and after gelation, they were placed on APTS functionalized coverslips (**Fig. 1D**) or directly transferred to any classical multi-well plate (**Fig. 1E**).

This method enabled us to generate hundreds of homogenous spheroids per microsystem in each well of a multi-well plate (**Fig. 1E, F**). Thanks to the hydrogel nature of the microwells, many experimental steps including rinsing, changing medium, spheroid fixation and immunostaining could be implemented in the same multi-well plate, with no manipulation of spheroids, which resulted in the treatment and labelling of several spheroids simultaneously (**Fig. 1G**). The advantage of the agarose microwells was the efficient transfer of medium and solutions through it. The exchange rate has been quantified by following-up the removal of FITC dye and AGuIX^®^-Cy5.5 nanoparticles from the agarose microwells via time-lapse image acquisition using confocal microscopy (**Fig.SI.3**). All curves were exponentially decreasing with a characteristic time of 25 min for FITC (23-27 min depending on the depth) and of 1 to 2 hours for AGuIX^®^-Cy5.5 nanoparticles, depending on the depth of the focal plane. A plateau is reached after two hours for FITC (at 25±5 %) and after 10 hours for AGuIX^®^-Cy5.5 nanoparticles (at 5±3 %).

Of note, the compatibility of the hydrogel-based microsystem with coverslips enabled *in-situ* quantification of nanoparticle penetration and their 3D distribution within spheroids with high-resolution optical microscopy such as confocal fluorescence microscopy (**Fig. 1H**). All spheroids were within the same focal plane, giving access to easy parallelization of 3D spheroids imaging (**Fig. 1I**). This is an important aspect compared to already proposed hydrogel microwells, where spheroids need to be transferred to a dedicated microscopic-plate for high-resolution 3D imaging ^27,49–51^. Such transfer first increases the complexity in terms of handling and imaging, and second may induces fusion between spheroids, or deformation of spheroids, which in turn may introduce biases in the analysis. Our original and simple process (Biocompatible hydrogel microwell plate, under patent^39^) bridge an important gap for in-depth optical spheroid analysis.

Moreover, these microwells are compatible with time-lapse optical microscopy, facilitating follow-up of spheroids growth for several days (**Fig. SI.4, movie 1**). The system enables us to produce very homogenous spheroids (**Fig. SI.5**), which gives access to the heterogeneity of cell response, with no bias induced by size heterogeneity. In our study, nanoparticles penetration was mainly evaluated using fluorescent intensity obtained from 3D confocal image acquisition. Taking advantage of the large statistics provided by our microsystems, we assessed the minimum number of spheroids required to get reliable results (**Fig. SI.6**). A minimum of N=30 spheroids is recommended to get reliable results at an imaging depth corresponding to the first quarter of the spheroids (0-50 μm from the periphery). This number rises to N=70 spheroids for an accurate analysis close to the equatorial plane.

### Cellular uptake of AGuIX^®^-Cy5.5 nanoparticles in 2D and 3D

As a proof-of-concept of the relevance of this new hydrogel-based microsystem, the penetration and distribution of AGuIX^®^-Cy5.5 nanoparticles (D_H_=5 nm) within spheroids was investigated using the colorectal cancer cell line HCT-116. In a previous study, it has been proven that localization of Gd-based nanoparticles tagged with Cy5.5 is the same as label-free nanoparticles in U87 cells ^52^. After 48 h of growth within 200-μm agarose microwells, spheroids were incubated with three different concentrations of AGuIX^®^-Cy5.5, 0.8, 1.5 and 2 mM in Gd, selected according to previous studies performed in 2D cell culture ^53^. In **Fig. 2A**. fluorescence images display the distribution of AGuIX^®^-Cy5.5 nanoparticles in spheroids after 24 h incubation with the three different concentrations at different depths. These images showed qualitatively that the number of nanoparticles clusters in spheroids directly increases with increase in initial concentration of AGuIX^®^-Cy5.5 nanoparticles. For 0.8 mM, very few nanoparticle clusters could be observed, while the number of clusters increased in 1.5 and 2 mM concentrations. For 2 mM concentration, nanoparticles were detected within the deeper layers of spheroids. Taking the spherical geometry of the sample into account to quantify the fluorescence in each images of spheroids, the mean intensity was calculated by averaging intensity along theta and phi angles in direction of the radius. The *in-situ* fluorescence analysis of **Fig. 2B** enabled to decipher the relative differences in nanoparticles penetration in the range of concentration analysed. Consistent with the fluorescent images in **Fig. 2A**, the mean intensity increased as the incubation concentration increased. From the outermost layer to the centre of the spheroids, the mean intensity decreased differently depending on the concentration (from 34±8% to 28±14% for 0.8 mM, from 68±8% to 46±15% for 1.5 mM and from 66±5% to 60±24% for 2 mM). For the largest concentration (2 mM), deep penetration was possible, while the penetration decreased exponentially with the depth for 1.5 mM, with a characteristic length of 44±2 μm. Such difference could be attributed to the higher number of nanoparticles reaching the centre of the spheroids for an incubation with 2 mM Gd. The relative independence of fluorescence intensity with depth for the lowest concentration (0.8 mM), could be attributed to a level close to noise, with no real penetration of nanoparticles nor in the periphery, neither in the centre of the spheroids.

**Figure.2.**
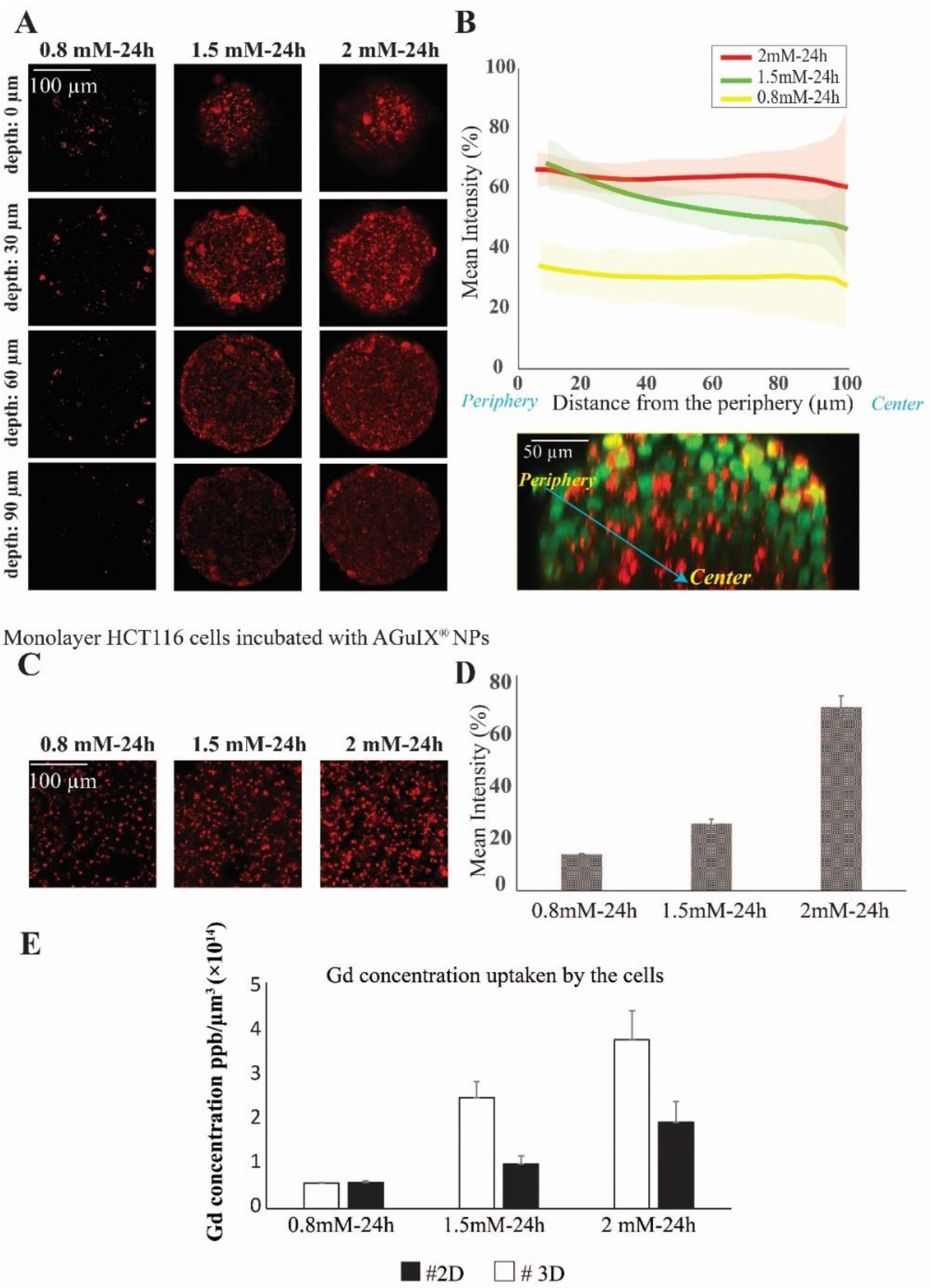
Quantification of penetration and cellular uptake of AGuIX^®^-Cy.5.5 nanoparticles in HCT116 tumour spheroids and monolayer cell culture. **(A)** Representative confocal fluorescence images of HCT-116 spheroids incubated with 0.8, 1.5 and 2 mM concentration of AGuIX^®^-Cy 5.5 for 24 h for four different depths (0, 30, 60 and 90 μm). **(B)** Mean intensity along with standard deviation (light colors) of AGuIX^®^-Cy 5.5 as a function of the distance from the spheroids periphery (see the orthogonal view in the inset, green=nuclei, red = AGuIX^®^-Cy5.5) for 0.8 mM (yellow, N=73), 1.5 mM (green, N=68) and 2 mM (red, N=121), three independent experiments. **(C)** Representative confocal fluorescence images of monolayer HCT-116 cells exposed to AGuIX^®^-Cy5.5 nanoparticles with 0.8, 1.5 and 2 mM. **(D)** Quantification of the mean intensity of AGuIX^®^ -Cy5.5 nanoparticles in maximal projection of confocal fluorescence images of monolayer cells after 24h incubation with different AGuIX^®^-Cy5.5 concentrations: 0.8 mM (yellow, N=40), 1.5 mM (green, N=40) and 2 mM (red, N=40), three independent experiments. Error bars represent the standard deviations. **(E)** Mean and standard deviation of the concentration of Gd (ppb/μm3) uptaken by the cells after incubation with 0.8, 1.5 and 2 mM concentration of AGuIX^®^ for 24 h in HCT116 cell spheroids and monolayer cell culture measured with ICP-MS technique (N = 6, two independent experiments).

To be sure that the presence of agarose in our microsystem does not affect the distribution and cellular uptake of AGuIX^®^-Cy5.5 nanoparticles within spheroids, a control experiment was made using ultra low adhesion 96-well plate and 2mM AGuIX^®^-Cy5.5 nanoparticle concentration **(Fig. SI 7)**. Similar results concerning the penetration of the nanoparticles were obtained: same normalised intensity range, and similar evolution as a function of distance from the periphery.

Deep penetration of small nanoparticles (<12 nm) within deep interstitial space have already been reported *in vivo* ^54^. The *in vitro* platform described in the current study enables to assess more quantitatively such penetration. It will hence be a valuable tool to relate such penetration with therapeutic efficacy in future studies.

To make a direct comparison with cellular uptake in 2D cell culture, monolayers of HCT-116 cells were incubated with the same concentrations of AGuIX^®^-Cy5.5 nanoparticles (**Fig. 2C**). As expected, the number of AGuIX^®^-Cy5.5 clusters raised as the initial concentration increased and the quantification of fluorescence images of cell monolayers (based on the mean intensity of AGuIX^®^-Cy5.5) confirmed that the uptake of nanoparticles increased with the concentration of AGuIX^®^-Cy5.5 in the incubation medium (**Fig. 2D** from 14.0 ±0.3 % for 0.8 mM, 25.8 ± 1.8 % for 1.5 mM to 70.8 ± 4.4 % for 2 mM). This mean intensity evolution was hence different from the one obtained in 3D in the periphery. However, as a true quantitative comparison is not possible using fluorescent analysis, elemental analysis by ICP-MS was performed concurrently to get a quantitative analysis of Gd content within cells for both 2D and 3D models (**Fig. 2E**). While the average nanoparticles uptake per cell in 2D and 3D were similar for 0.8 mM ((0.580±0.006).10^−14^ ppb/μm^3^ in 2D vs (0.59±0.05).10^−14^ in 3D), the uptake was two-fold higher in 2D compared to 3D for both 1.5 mM ((2.5±0.5).10^−14^ ppb/μm^3^ in 2D vs (1.0±0.2).10^−14^ in 3D) and 2 mM ((3.8±0.9).10^−14^ ppb/μm^3^ in 2D vs (1.9±0.6).10^−14^ ppb/μm^3^ in 3D).

One of the reasons of the reduction in effectiveness of therapeutics *in vivo* compared to monolayer cell cultures is the lack of efficient penetration and distribution of therapeutics throughout tumour tissue ^55^. This is what we also observed here, with a large reduction of nanoparticles uptake in 3D compared to 2D cell-culture.

Another approach was used to compare cellular uptake of AGuIX^®^-Cy5.5 nanoparticles in 2D and 3D: 2D cells were treated with 2mM AGuIX^®^-Cy5.5 nanoparticles for 24 h, then spheroids were made from these AGuIX^®^-Cy5.5 labelled cells using the usual protocol **(Fig SI 8)**. Interestingly, the distribution of nanoparticles differs when spheroids are made with already labelled cells, compare to direct incubation with already formed spheroids, further highlighting the difference in nanoparticles availability between 2D and 3D models.

### Kinetics of AGuIX^®^-Cy5.5 nanoparticles transport into spheroids

One of the crucial parameters in nanoscale design is the pharmacokinetics of nanoparticles and understanding this aspect of cell-nanoparticles interactions has a great importance ^56,57^. The kinetics of penetration of AGuIX^®^-Cy5.5 nanoparticles within HCT-116 cell spheroids grown for 48 h were assessed by analysing confocal images obtained for different incubation times (1, 24 and 72 h), for the highest concentration investigated (2 mM) (**Fig. 3A**). After 1 h incubation, the AGuIX^®^-Cy5.5 nanoparticles were mostly residing in the peripheral layer of the spheroids, especially in the extracellular space. After 24 h, clusters of nanoparticles were found throughout the spheroids. At 72 h, the number of clusters were increasing for all depths, up to the equatorial plane (100 μm).

**Figure.3.**
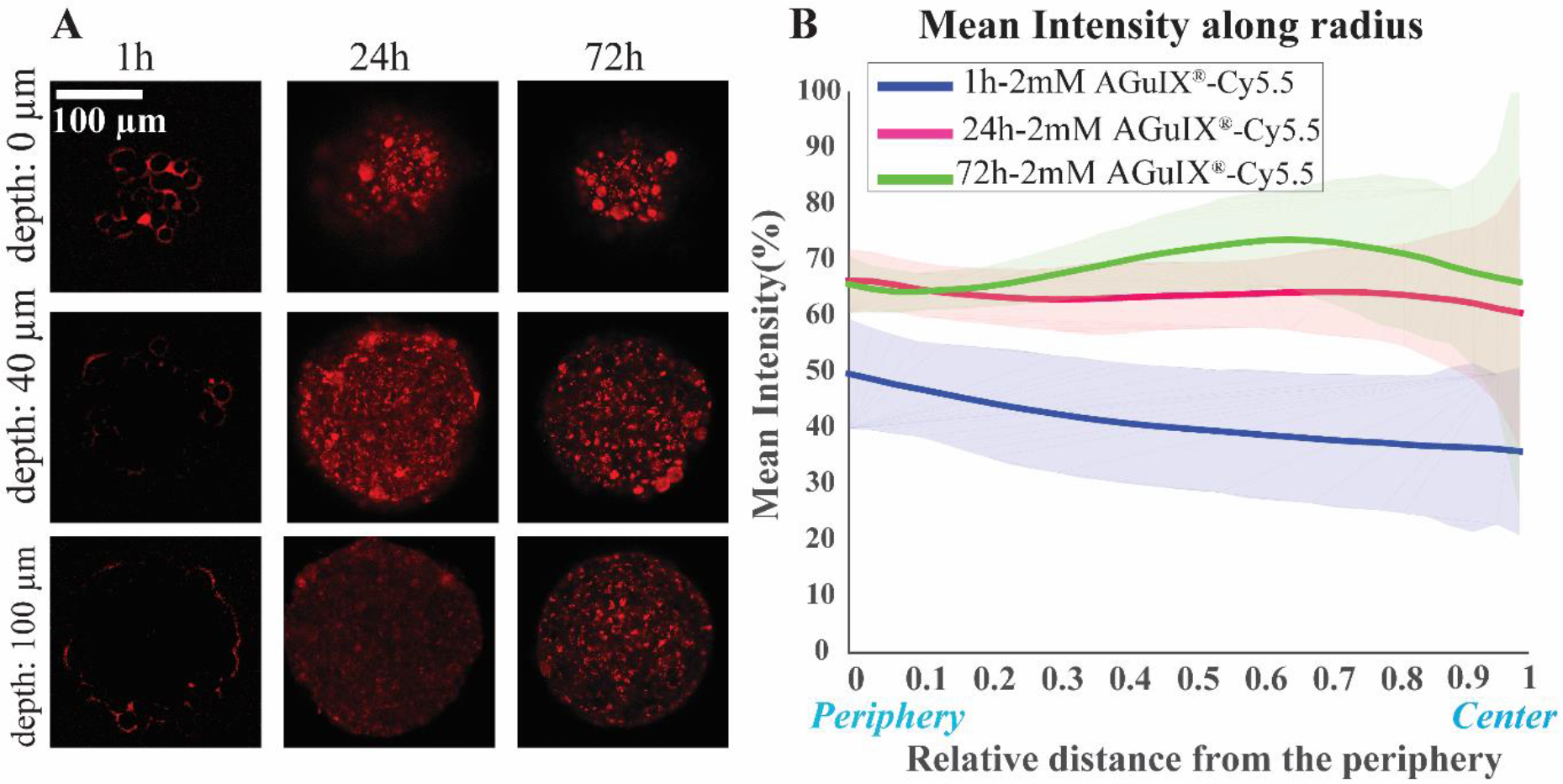
Kinetics of penetration of AGuIX^®^-Cy5.5 nanoparticles in HCT-116 cell spheroids. **(A)** Representative confocal fluorescence images of HCT-116 grown for 48 h, exposed to 2mM concentration of AGuIX^®^-Cy5.5 nanoparticles for 1, 24 and 72 h. **(B)** Mean intensity along with standard deviation (light colours) as a function of relative distance from the periphery for 1 h (blue, N=50), 24 h (magenta, N=121) and 72 h (green, N=63), three independent experiments.

The average intensity was exhibiting different evolution with the distance from the periphery, depending on the incubation time (**Fig. 3B**). At the periphery, the average intensity was lower for 1 h incubation (52.2±1.3%) than for 24 h and 72 h that exhibit similar values (68±1% and 65.8±0.7%, respectively). When we moved to the centre of the spheroids, the mean intensity is slightly lowered for 1 h and 24 h incubation (from 52.2±1.3% down to 38.7±2.7% for 1h, from 68±1% down to 59.5±1.9 % for 24 h). Accordingly, in 1 h incubation the mean intensity is less than 24 h and 72 h samples in all regions of the spheroids. Surprisingly, for 72 h, the average intensity exhibited a non-monotonous evolution with the distance from the periphery, with an intensity larger for middle layers than at the periphery (66±5% vs 73.8 ±8.5% at a depth of 60 μm). This may be the results of increased number of clusters found for intermediate layers after 72 h incubation time.

To follow the distribution and the transport of nanoparticles within spheroids, an experiment was designed to assess changes in AGuIX^®^-Cy5.5 nanoparticles distribution before and after rinsing steps (**Fig. 4**). In this experiment, 2 days after cell seeding (**Fig. 4 step I**), the formed HCT-116 spheroids were incubated with 2 mM AGuIX^®^-Cy5.5 solution for 72 h (**Fig. 4, step II**). Spheroids were then imaged in incubation medium (**Fig. 4 step III**) and after three washing steps of 15 min each (**Fig. 4 step IV**). Spheroids were kept in the incubator for additional 24 h and then imaged before (**Fig. 4 step V)** and after (**Fig. 4 step VI**) another washing procedure. Confocal fluorescence microscopy of living spheroids showed that AGuIX^®^-Cy5.5 nanoparticles fluorescence signals of the surrounding background (fluorescence signal outside spheroids) was decreasing gradually with the different washing steps for all depth (**Fig. 4A-D**). This is confirmed by the quantification of the mean intensity along the spheroid radius (**Fig. 4E**): while the mean intensity before washing (red curve) at the periphery was around 92±10%, it was decreasing to 59±10% at 55 μm distance from the periphery. After the first washing step (yellow curve), the mean intensity at the periphery reduced to 66±6% and reached a similar intensity level than to the one before washing at 55 μm distance from the periphery (58±4%). Further washing steps further reduced the mean intensity at the periphery (64±14% and 54±14% before and after the second washing step), while the mean intensity obtained for deeper layers exhibited similar levels. The second washing (blue curve), led to a steady value of mean intensity (~54%) close to the mean intensity obtained at 55 μm distance from the periphery of spheroids for all washing steps. This mean intensity should correspond to the signal coming from nanoparticles that are internalized by the cells, as all nanoparticles residing in the extracellular space have been washed away.

**Figure.4.**
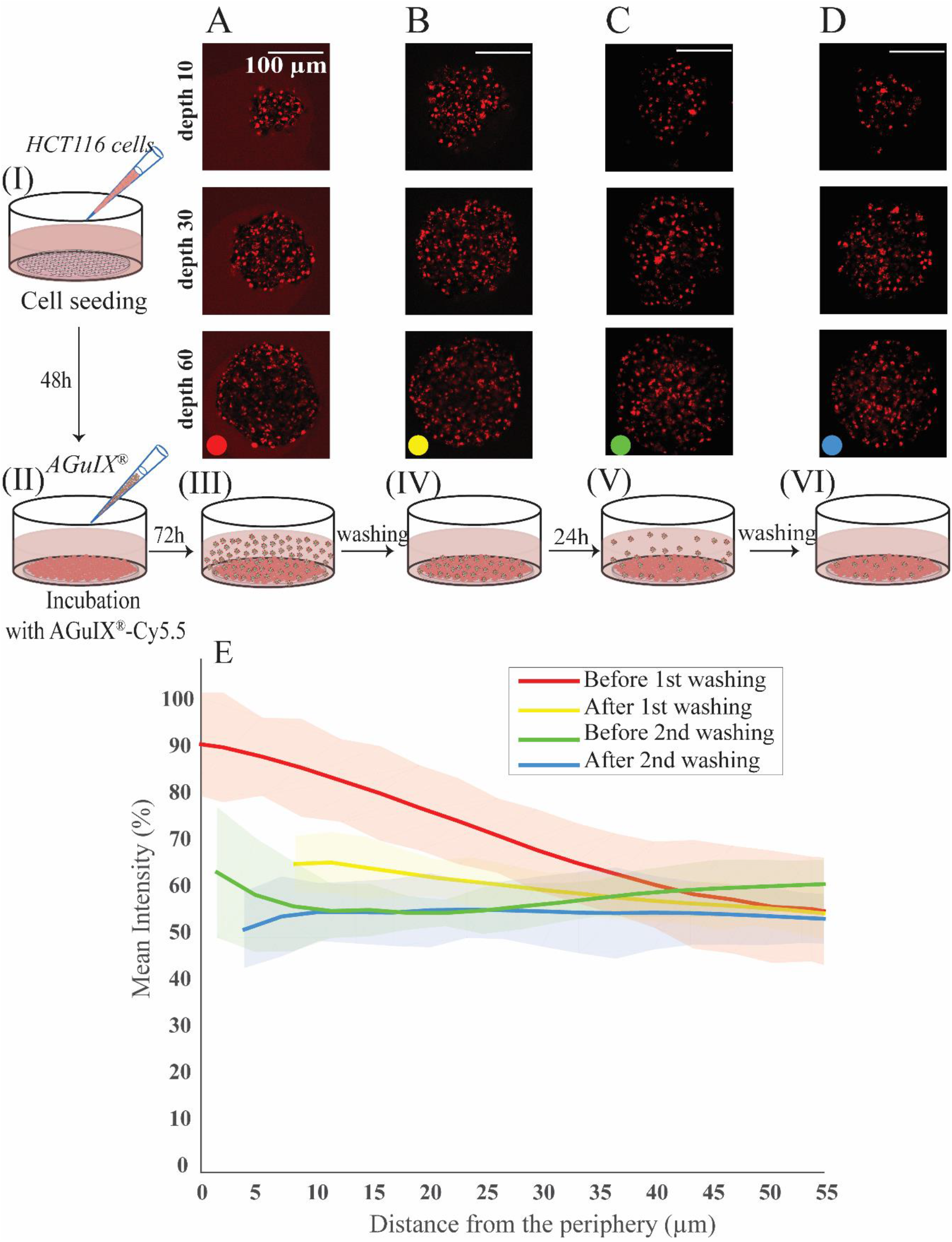
Dynamic analysis of AGuIX^®^-Cy5.5 nanoparticles transport and localization within spheroids. HCT-116 cell spheroids were prepared using agarose microsystem **(step I).** After 48 h growth, they were exposed to 2 mM AGuIX^®^-Cy5.5 solution for 72 h **(step II)**. **(A)** Spheroids were imaged in the incubation medium **(step III)**. **(B)** Spheroids were then rinsed with fresh medium three times for 15 min each and were imaged again **(step IV). (C)** Spheroids were allowed to grow for an additional 24h (in an incubator at 37°C and 5% CO_2_) before imaging **(step V). (D)** Spheroids were rinsed again with fresh medium (3×15min), before imaging **(step VI)**. **(E)** Quantification of AGuIX^®^-Cy5.5 nanoparticles mean intensity along the distance from the periphery (N=25). Bold lines represent the mean intensities, averaged for all spheroids. Light colours represent the standard deviations.

### Localization of AGuIX^®^-Cy5.5 nanoparticles in spheroids using their chemical signature

Due to the limit of resolution using standard confocal optical microscopy (200 nm in the best imaging conditions), only clusters of nanoparticles can be detected. In addition, we cannot rule out that the distribution of the fluorophore do not truly represent the distribution of the nanoparticles themselves. To confirm the presence of nanoparticles, nanoscale Secondary Ion Mass Spectrometry (nanoSIMS) was performed on spheroids sections (**Fig. 5**). This analytical technique allows the acquisition of elemental composition maps with a spatial resolution down to 50 nm. The images of ^12^C^−^ (see **Fig.SI.7**) and ^35^Cl^−^ (data not shown) indicates the absence of defect in the sample section. Any damage, even tiny holes, would appear with high contrast in signal, and such signal was not observed. This validated that the signal measured was originated from the sample and not from the subjacent pure silicon substrate. The image of ^12^C^14^N^−^ showed the histological aspect of the cell (data not shown) while the one of ^31^P^−^ (**Fig. 5A**) highlights the cell nucleus. Since AGuIX^®^-Cy5.5 nanoparticles are mainly made of Si, the images of ^28^Si^−^ allowed the observation of the chemical signature of the nanoparticles (**Fig. 5B**). Thereby, the nanoparticles were found unequivocally inside the spheroid, exclusively in the cytoplasm of the cells. Of note, again our microsystems enabled an easy sample preparation, as all spheroids were within the same sectioning plane.

**Figure 5.**
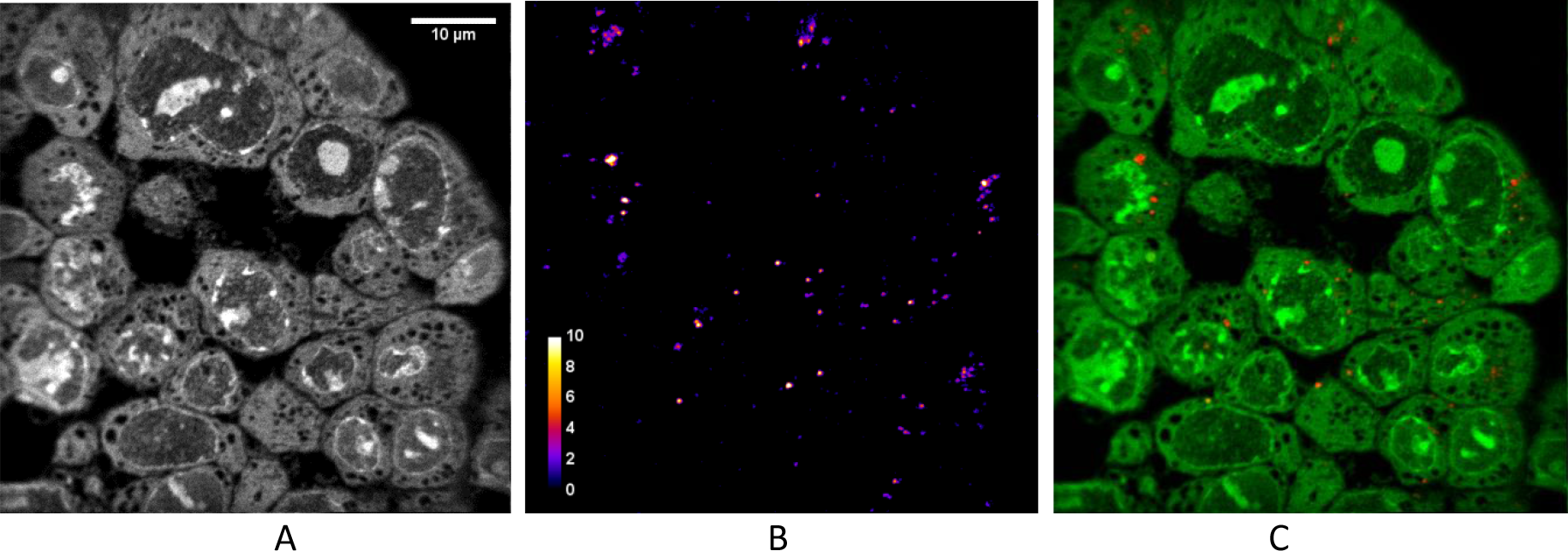
Localisation of the AGuIX^®^-Cy5.5 nanoparticles in spheroids using NanoSIMS. NanoSIMS images of HCT-116 spheroids loaded with AGuIX^®^-Cy5.5 nanoparticles. **(A)** corresponds to the signal of ^31^P^−^ showing the cell structure. **(B)** highlights the signal of ^28^Si^−^ representing the intracellular location of AGuIX^®^-Cy5.5 nanoparticles. **(C)** Merged image of ^28^Si^−^ and ^31^P^−^. Scale bar: 10 μm.

### Localisation of AGuIX^®^-Cy5.5 within cells in 2D and 3D using immunostaining

Thanks to the full compatibility of the microsystems with *in-situ* immunostaining, it was possible to assess the localisation of nanoparticles in 2D cells (**Fig. 6**) and multicellular tumour spheroids (**Fig. 7**), using confocal fluorescence microscopy.

**Figure 6.**
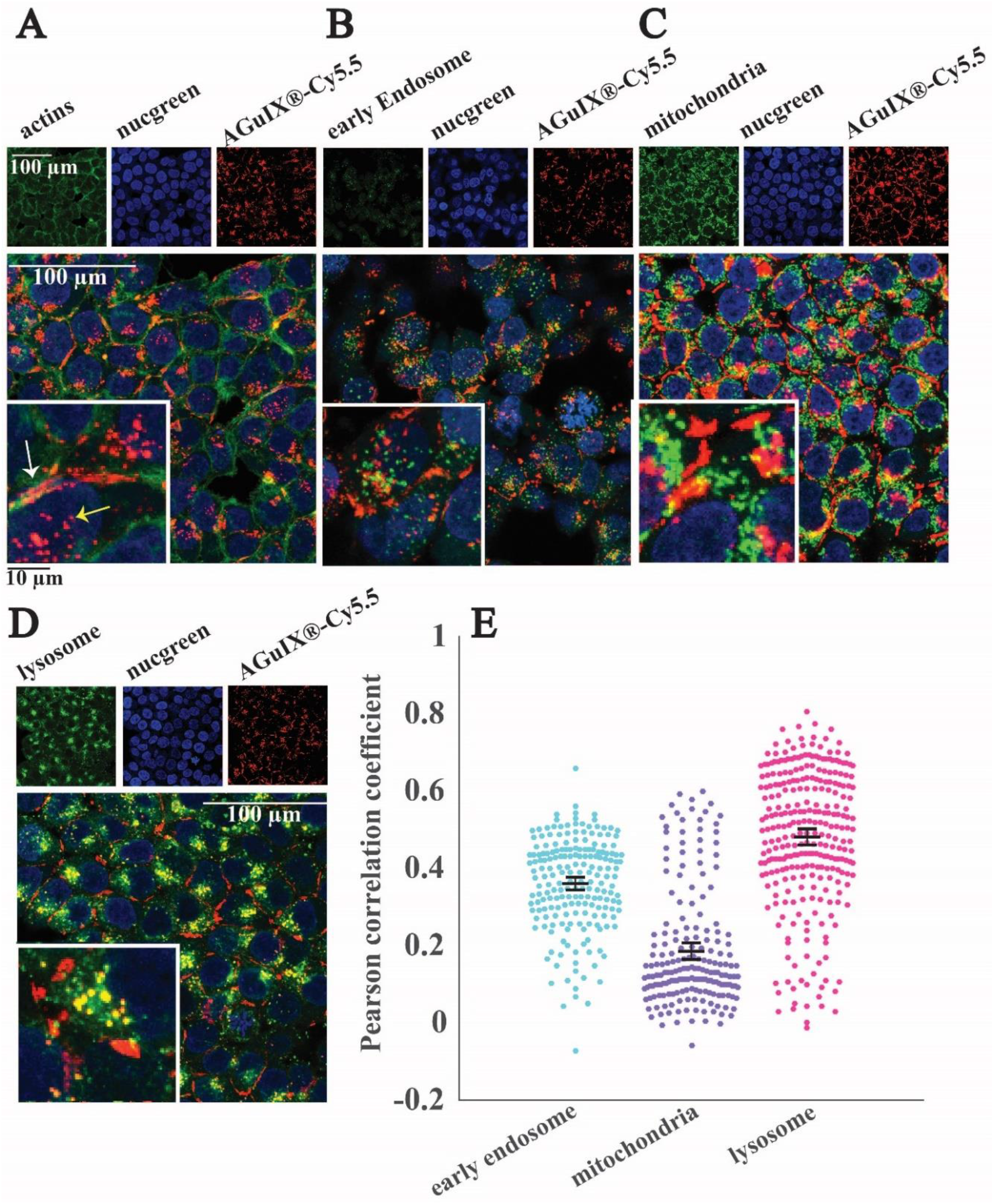
Localization of the AGuIX^®^-Cy5.5 nanoparticles in 2D monolayers. Fluorescence images of HCT-116 cells incubated with AGuIX^®^-Cy5.5 nanoparticles (2 mM, 24 h) and immunostained with antibodies to find colocalization of nanoparticles inside cells. In all images red channel and blue channel represent AGuIX^®^-Cy5.5 nanoparticles and cell nuclei respectively. **(A)** Green channel depicts phalloidin, a marker of actin in cells, which demonstrates nanoparticles localising both inside cells (yellow arrow) and in the space between cells (white arrow). **(B)** Green channel shows early endosome in the cells, with no colocalization with AGuIX^®^-Cy5.5 nanoparticles. **(C)** Green channel shows mitochondria and reveal very low colocalization with AGuIX^®^-Cy5.5 nanoparticles (yellow colour). **(D)** Green channel shows the lysosomes and colocalization is demontrasted by the yellow colour. White scale bar, 100 μm and black scale bar, 10 μm. **(E)** Distribution of Pearson correlation coefficients in the different fields of view to quantify the colocalization of AGuIX^®^-Cy5.5 nanoparticles with the three different cell organelles investigated. Error bars represent standard error of the mean (SEM) of pearson correlation coefficient values obtained for all fields of view and all available depth, for three independent experiments. It was plotted as scatter plots using the Matlab UnivarScatter function (^©^Manuel Lera Ramírez, 2015, available in matlab exchange files).

**Figure 7.**
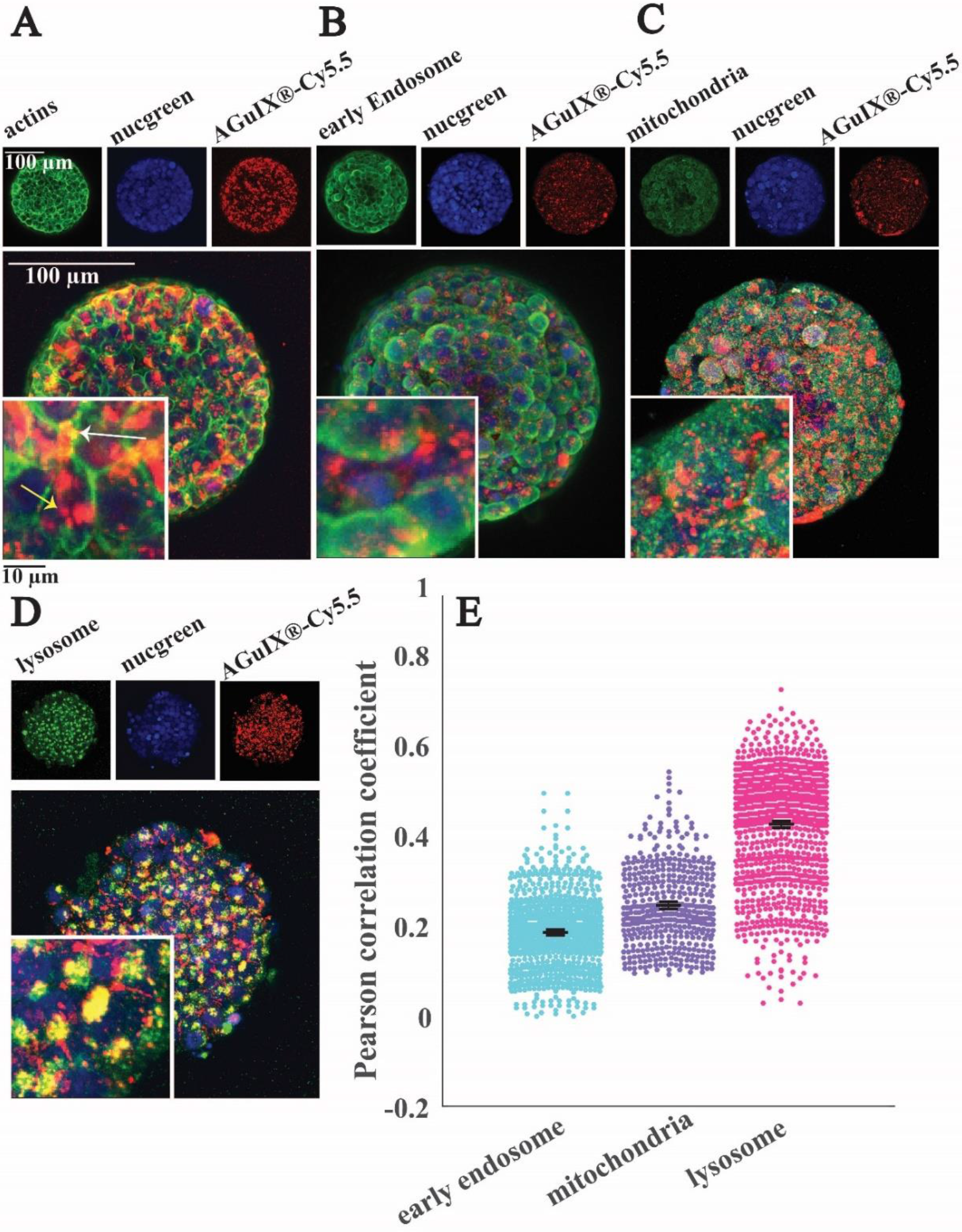
Localization of AGuIX^®^-Cy5.5 nanoparticles within spheroids. Fluorescence images of HCT116 cells spheroids incubated with AGuIX^®^-Cy5.5 nanoparticles (2 mM, 24 h) and immunostained with antibodies to find colocalization of nanoparticles inside cells. In all images, red and blue channels represent AGuIX^®^-Cy5.5 nanoparticles and cell nuclei respectively. **(A)** Green channel depicts phalloidin, a marker of actin in cells, which demonstrates that nanoparticles localize both inside cells (yellow arrow) and in the extracellular space of spheroid (white arrow). **(B)** EEA1 antibody in green channel shows early endosome; with very low colocalization with nanoparticles (yellow colour)**. (C)** AIF antibody labelled mitochondria is shown in green, with very low colocalization with AGuIX^®^-Cy5.5 nanoparticles (yellow colour). **(D)** LAMP1 antibody in green channel stains lysosomes. Yellow colour represents the colocalization of nanoparticles in red and lysosomes in green. White scale bar, 100 μm and black scale bar, 10 μm. **(E)** Quantification of the colocalization of AGuIX^®^-Cy5.5 nanoparticles with the three different cell organelles investigated using Pearson correlation coefficient. Distribution obtained for all imaged spheroids and all imaging depth for the three different cell organelles. Error bars represent standard error of the mean (SEM) of pearson correlation coefficient values obtained for all fields of view and all available depth, for three independent experiments. It was plotted as scatter plots using the matlab UnivarScatter function (^©^Manuel Lera Ramírez, 2015, available in Matlab exchange files).

Labelling of cell organelles confirmed that nanoparticles were present in both extracellular and intracellular space in 2D cells (**Fig. 6A**) and in 3D spheroids (**Fig. 7A**). Very low colocalization of AGuIX^®^-Cy5.5 with early endosomes (**Fig. 6B, E in 2D and Fig. 7B, E in 3D**) or mitochondria (**Fig. 6C, E in 2D and Fig. 7C, E in 3D**) was evidenced by immunostaining, while a large colocalization with lysosome was observed in both 2D (**Fig. 6D, E)** and 3D environments (**Fig. 7D, E**). Pearson correlation coefficient in both 2D cells and 3D spheroids (**Fig.6E** in 2D and **Fig.7E** in 3D) showed a higher value for lysosomes (0.48±0.18 and 0.42±0.12 for 2D and 3D respectively, compared to 0.36±0.12 and 0.18±0.09 for early endosomes in 2D and 3D and 0.19±0.15 and 0.24±0.09 for mitochondria in 2D and 3D) showing the main intracellular localisation of AGuIX^®^-Cy5.5 nanoparticles. Noteworthy, this colocalization was not total and some nanoparticles were still residing in between cells. These outcomes are in accordance with previous studies showing localisation of nanoparticles in endocytic pathway and in lysosomes^38,52^. Internalization mechanisms of AGuIX^®^ have been thoroughly investigated in 2D ^38^. It has been shown that entry of such sub-5nm nanoparticles is different depending on nanoparticles concentration: passive diffusion and eventually macropinocytosis, in case of formation of nanoparticles cluster at the surface of the cell. It is known that the internalization pathway for a specific nanoparticle can differ between cell lines ^58^. For the HCT-116 cell line used in this study, localisation of AGuIX^®^-Cy5.5 nanoparticles in lysosomes and in smaller amounts in early endosomes confirms they were likely internalized by an endocytic mechanism ^59^. Despite dominant colocalization for both 2D and 3D with lysosomes, in 2D images the Pearson Correlation Coefficient average value for early endosomes is higher than mitochondria **(Fig. 6B, E)**, which contrasts with these values in 3D **(Fig. 7B, E).** One explanation for this difference is that for spheroids, cells have varying access to the nanoparticles depending on their spatial position within spheroids, which could lead to different internalization processes. In 3D spheroids, AGuIX^®^-Cy5.5 nanoparticles were confronted to barriers to reach the cells in deeper layers, therefore they reach deeper layers in a lower amount **(Fig.2B)** and with a delay **(Fig.3B)**, which can change their intracellular fate. This is another argument in favour of 3D system for nanoparticle transport analysis. In 2D, all cells are submitted to the same homogeneous concentration of nanoparticles, while in 3D, there is a large difference in nanoparticles availability in between cells that are at the periphery and cells in the centre of the spheroids. In addition, in spheroids, similar to natural tumours there is a gradient of pH, oxygen and metabolites ^60^, which might affect internalization and intracellular trafficking of nanoparticles in deeper layers ^61^.

As highlighted with the overall mean intensity decrease with washing procedure for the peripheral layers of the spheroids (**Fig. 4**), the extracellular nanoparticles were efficiently washed away after a long washing procedure (**Sup. Fig. SI 10**, no extracellular nanoparticles were detected with immunostaining). Similar to results obtained after 72 h incubation, colocalization with lysosomes was still the major localisation of nanoparticles after this extensive washing procedure (**Fig. 8 A, B, C for lysosomes compared to Fig. 8 D, E, F for mitochondria**). The comparison of Pearson’s correlation coefficient of AGuIX^®^-Cy5.5 with both lysosomes and mitochondria before and after washing suggests minor intracellular trafficking and/or exocytosis of nanoparticles over time (**Fig. 8B for lysosomes and 8E for mitochondria**). The Pearson’s correlation coefficient of AGuIX^®^-Cy5.5 with lysosomes remained within a similar range before and after washing (mean values of 0.41 ± 0.03 vs 0.40 ± 0.11 respectively), while a decrease in Pearson’s correlation coefficient of AGuIX^®^-Cy5.5 with mitochondria is observed after washing, particularly in outer layers (0.26 ± 0.02 vs 0.18 ± 0.04 at 10 μm-depth before and after washing respectively), reaching very low value for inner layers (0.20 ± 0.04 vs 0.18 ± 0.02 at 60 μm-depth before and after washing respectively). Such decrease could be attributed to the removal of few AGuIX^®^-Cy5.5 clusters residing in mitochondria or possible intracellular trafficking during the washing procedure. Hence, we could say that washing procedure had lesser effect on AGuIX^®^-Cy5.5 nanoparticles residing in lysosomes.

**Figure 8.**
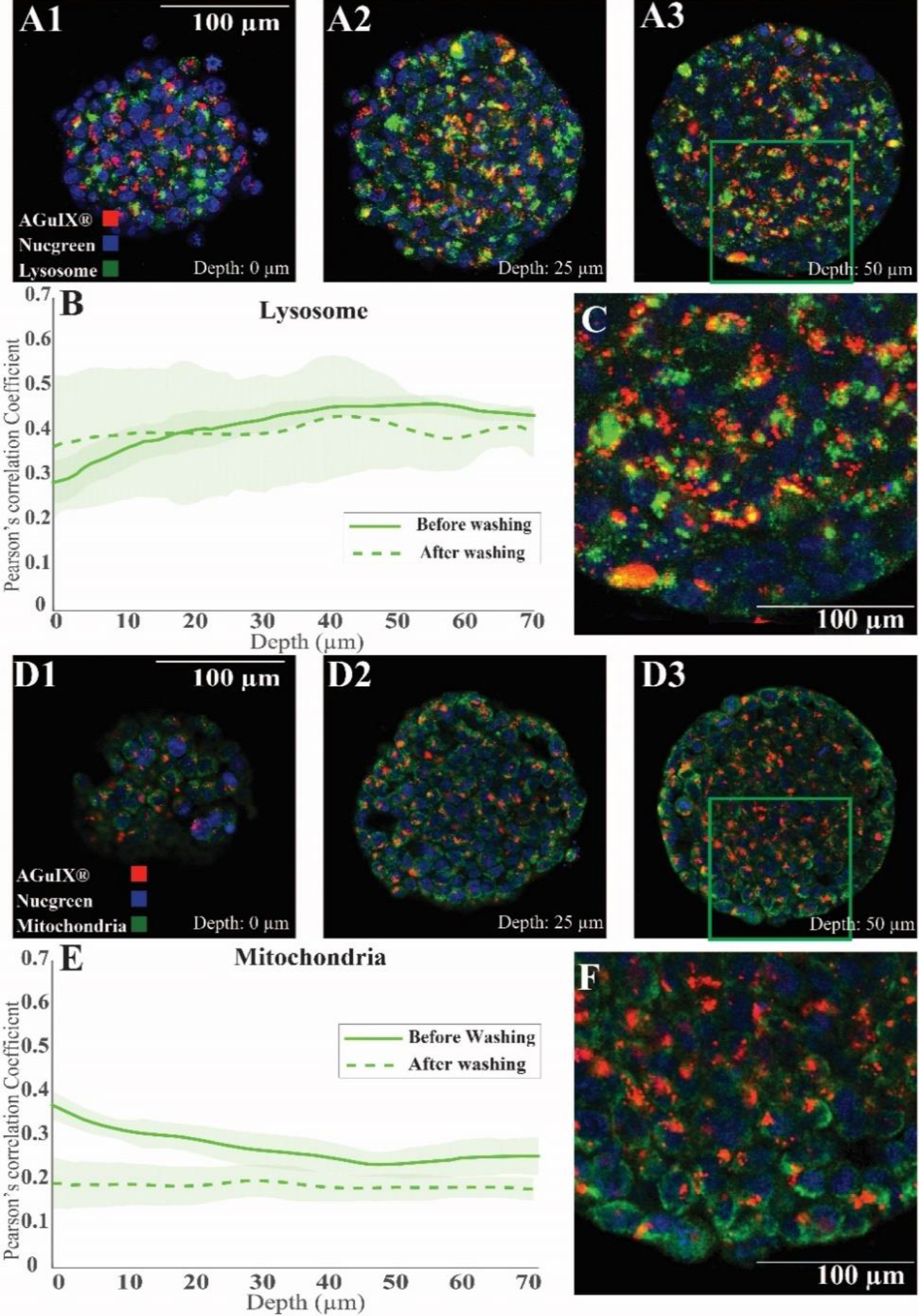
Localisation of Aguix-Cy5.5-nanoparticles after an extensive washing procedure. Confocal fluorescence images of HCT-116 spheroids incubated with AGuIX^®^-Cy5.5 nanoparticles for 72 h with 2 mM AGuIX^®^ solution, and washed according to the procedure mentioned in Figure 4, then fixed and immunostained with antibodies to find colocalization of nanoparticles in spheroids. For all images, red and blue channels are staining AGuIX^®^-Cy5.5 and nuclei respectively. **(A1-A3)** Representative images of lysosomes immunostaining obtained at various depth (**A1-0 μm, A2-25 μm, A3-50 μm**. Green channel = lysosome [LAMP1 antibody], yellow colour = possible co-localization of AGuIX^®^-Cy5.5 nanoparticles with lysosomes) **(B)** Pearson’s correlation coefficient for AGuIX^®^-Cy5.5 nanoparticles with lysosomes along with standard error of the mean (light colour) as a function of depth (n = 27 spheroids, 3 independent experiments before washing, n = 5 spheroids after washing). **(C)** Zoomed-in portion of merged image at a depth of 50 μm (square in A3). **(D1-D3)** Representative images of mitochondria immunostaining obtained at various depth (**D1-0 μm, D2-25 μm, D3-50 μm**. green channel = mitochondria [AIF antibody], yellow colour = possible co-localization of AGuIX^®^-Cy5 nanoparticles with mitochondria). **(E)** Pearson’s correlation coefficient for AGuIX^®^-Cy5.5 nanoparticles and mitochondria along with standard error of the mean (light colour) as a function of depth. (n=22 spheroids, 3 independent experiments before washing, n=5 spheroids after washing). **(F)** Zoomed-in portion of merged image at a depth of 50 μm (square in D3).

## Conclusion and outlooks

We show in this study a simple agarose-based microsystem to quantitatively track nanoparticles penetration and subcellular localisation within 3D-cell culture model. The reproducibility of the spheroid size obtained with such procedure dispenses the use of sophisticated automatic procedure to choose and pick the appropriate spheroids. Of note, our microsystems can be manufactured on conventional multi-well plates. It is hence fully compatible with available multi-well automated strategies^62^. In the present study, the proof-of-concept was validated using spheroids made with the classical colorectal cell line HCT-116. Nevertheless, our approach is fully compatible with primary cells from patients that could be grown as organoids ^63^ in our microsystems, combining full optical microscopy compatibility, size and shape reproducibility, and large statistics. Combined with optical and digital clearing ^64^, our approach opens up the possibility to resolve tumour heterogeneity, at the single cell-level, in a physiological context.

In the present study, the standard agarose used for the preparation of the microsystems provides a cell-repellent surface, with a stiffness in the 150 kPa range ^65,66^. In future studies, the mechanical properties of the agarose gels will be adjusted using different concentrations and type of agarose. Elastic hydrogels as soft as 1 kPa can be obtained using low concentration of ultra-low agarose ^65^, matching the physiological range of stiffness. It now calls for dedicated studies to assess how nanoparticles penetration and therapeutic efficacy is affected by the size of the 3D-cell assembly, the presence of an extracellular matrix of different stiffness and composition, and the presence of associated tumour cells ^67^.

## Acknowledgments

This work was supported by the “Institut Universitaire de France” (IUF). We thank L. Fuoco for her help in the early development of this project, R. Fulcrand for his support in photolithography, M.G. Blanchin for her help in spheroid preparation protocols for Electron Microscopy, as well as the CTμ platform (Centre Technologique des Microstructures) for resin embedding of spheroids.

## Supplementary Figures

### Figure SI 1: Quantification of photon penetration within spheroids

In optical imaging of thick three-dimensional biological samples, light scattering due to mismatch of refractive index between cellular components limits imaging of deep layers in these samples ^68^. To deal with this issue and to evaluate clarification techniques on enhancement of confocal microscopy of multicellular tumour spheroids, HCT-116 spheroids were prepared via agarose microwells and fixed 72 h after cell seeding. Nuclei were then stained using Nucgreen™ -Dead 488. Two of the samples were incubated in RapiClear or 80%/20% glycerol/PBS solution overnight and the third sample was kept in PBS. All samples were then mounted in iSpacers (2×0.5 mm) with fresh clarification solutions or fresh PBS for control sample. Ten spheroids from each sample were imaged. A qualitative analysis of the images (**Fig. SI 1A**) shows that nuclei fluorescence signal is detected much deeper for clarified spheroids compared to unclarified ones. The orthogonal views of spheroids confirm this (**Fig. SI 1B**). These images were analysed using a routine prepared in Matlab to measure the average fluorescence intensity along the spheroids radius (**Fig. SI 1C**). The mean Intensity of spheroids clarified with glycerol (Green curve) shows the highest mean intensity for all regions in spheroids compared to the two other samples. While the mean intensity of RapiClear-clarified spheroids (Red curve) is lower than glycerol-clarified spheroids in all regions of the spheroids, the intensity decay is similar for both clarified solutions. The 80% glycerol solution was hence selected as the standard clarification technique for all this study.

### Figure SI 2: Detailed mounting procedure for imaging

There are different possibilities to mount the microwells for optical imaging, depending if one wants to acquire fixed, live spheroids, or to follow spheroids over time using time-lapse.

1-For fixed spheroids, the easiest and quickest way is to mount the microwells between 2 coverslips, separated by a 1 mm sticky spacer (two 0.5 mm-thick iSpacer provided by SunJin Lab were used for their convenience, but other spacers could also be used, **Fig. SI2 B**).

2-For live spheroids, it is possible to transfer the microwells in optical imaging chamber (such as Ibidi^®^ 8-well plate, **Fig. SI2 C**).). We used such possibility in preliminary experiments to check that distribution of nanoparticles were not modified by fixation procedure (data not shown), as well as for the assessment of nanoparticles transport and localization after extensive washing procedure (**Fig. 4**).

3-For time-lapse follow-up, to avoid any drift during acquisition, it is necessary to directly bond the microwells to the coverslips. This is possible using APTS-functionalized coverslips (representation in **Fig. 1D**, patented process^39^). We used such procedure for growth monitoring using time lapse microscopy (**Fig. SI 4**).

### Figure SI 3: Quantification of the removal of FITC and AGuIX^®^-Cy5.5 nanoparticles in agarose-based microwells

To understand the ability of agarose gel in transporting molecules, the agarose-based microsystems were incubated with either FITC solution (0.05 mM in PBS), either AGuIX^®^-Cy5.5 nanoparticles (2 mM in complete culture medium). The fluorescent solution was then replaced with PBS (for FITC) or culture medium (for AGuIX^®^-Cy5.5) and the decrease in fluorescence intensity was followed by time-lapse confocal microscopy. The images of different depths of agarose-based microwells were analysed using a Matlab routine quantifying the mean intensity changes over time (**Fig. SI 3**). For FITC (**Fig.SI3 A**), after the first two hours, there is a 75±5% reduction in the initial mean intensity in the microsystem, reaching a plateau at 25±5 % depending on the depth of the focal plane. All curves were exponentially decreasing with a characteristic time of 25 min (23-27 min depending on the depth of the focal plane). As expected, as AGuiX^®^-Cy5.5 nanoparticles (D_H_=5 nm) are much larger than FITC (Mw=376 g/mol, D_H_~0,25 nm), the diffusion is one order of magnitude slower than with FITC, but still efficient in the agarose, both when culture medium is replaced by AGuIX^®^-Cy5.5 (**Fig.SI3 B)**, or when AGuIX^®^-Cy5.5 is replaced by culture medium (**Fig.SI3 C).** When culture medium is replaced by AGuIX^®^-Cy5.5, the characteristic time obtained for the deepest part of the gel is of the order of 22-24 min (23,9 ±0.4 min for Z=0, 23,6 ±0.6 min for Z=80 μm and 22,4 ±1,7 min for Z=160 μm), while it already reached the maximum intensity upon imaging for the upper part (z=240 μm). When AGuIX^®^-Cy5.5 is replaced by culture medium (after 3×15 min washing, following the procedure done for all experiments), the fluorescent is decreasing with a characteristic time of the order of 1-2h, depending of the depth (69 ±5 min for Z=0, 82±6 min for Z=80 μm, 104 ±5 min for Z=160 μm and 129 ±5 min for Z=240 μm).

### Figure SI 4. Time-lapse follow-up of spheroid growth

To understand the influence of AGuIX^®^-Cy5.5. nanoparticles on the cell proliferation and growth rate of HCT-116 cell spheroids, cells were seeded in agarose-based microwells using APTS-functionnalised coverslips. This procedure enables live imaging with no drift of the microwells over-time. After 48 h, the HCT-116 spheroids were exposed to AGuIX^®^-Cy5.5 nanoparticles with three different concentrations (0.8, 1.5 and 2 mM). Control samples with no AGuIX^®^-Cy5.5 nanoparticles were also monitored in parallel. The growth of spheroids was followed by time-lapse optical microscopy during three days of incubation with AGuIX^®^-Cy5.5 nanoparticles (time interval between each image acquisition = 4 h). These images were manually segmented using a dedicated routine in Matlab. Then, from the projected area, an equivalent diameter was computed, and making the assumption of spherical shape, the equivalent volume of each spheroids was calculated. Spheroids growth is followed by representing the relative evolution of the volume over time (Volume normalised by the initial volume [at day 2]).

### Figure SI 5. Characterization of spheroids size distribution from Day 2 to Day 4 after cell seeding

One advantage of using agarose-based microwells to prepare multicellular tumour spheroids is the homogeneity of spheroids size. To show the homogeneity of spheroids, the equivalent diameter for each spheroid was calculated during the growth follow-up **(Fig. SI 4)**. From this, the distribution of spheroids diameter in day 2, day 3 and day 4 for control conditions were plotted using the UnivarScatter matlab function developed by Manuel Lera Ramírez (Copyright (c) 2015).

### Figure SI 6. Importance of statistics

Due to cell heterogeneity, large statistical variances are expected, even if our process enables to generate very reproducible spheroids in terms of size. The variance on the Mean Intensity of the fluorescence signal of AGuIX^®^-Cy5.5 nanoparticles is analysed in **Fig.SI 6**, for different distance from the periphery.

The standard deviation of the normalized Intensity of AGuIX^®^-Cy5.5 obtained, is calculated as a function of the number (N) of spheroids, with a random sampling of N spheroids over the 121 spheroids acquired for this experimental condition (24h incubation with 2mM AGuIX^®^-Cy5.5). The random sampling is repeated 10 times to simulate 10 different experiments, and the mean of the obtained SD computed. The obtained SD first increases, until reaching a plateau around N=20-40 spheroids (**Fig. SI6, Inset**). The initial rising may be attributed to the heterogeneity among spheroids.

The plateau of the SD is increasing with the distance from the periphery (with a plateau at ~5% for 30 and 50 μm from the periphery, and up to ~10% for 80 μm from the periphery).

Once the plateau is reached, the standard error of the mean (SEM) and the relative standard errors (RSE=SEM/mean) on the normalized Intensity are therefore decreasing with N as N ^−1/2^ (**Fig. SI 6**). To get a RSE below 2%, N=20-25 spheroids are necessary for an analysis up to 60 μm from the periphery. For deep layers, a larger number of spheroids are needed to reach such RSE (N=70 spheroids for 80 μm from the periphery). Hence a minimum of N=30 spheroids is recommended to get reliable results at an imaging depth corresponding to the first quarter of the spheroids. This number rises up to N=70 spheroids for an accurate analysis close to the equatorial plane.

### Figure SI 7. Control experiments using Ultra-Low Adhesion multi-well plate

The goal of this experiment was to compare the distribution of AGuIX^®^-Cy5.5 nanoparticles in HCT-116 spheroids prepared in a traditional ultra-low adhesion 96-well plate with spheroids made in agarose microwells. HCT-116 cells were seeded at a density of 10 cells/well in a 96-well plate (200 μl culture medium per well), and spheroids were formed through self-assembly aggregation. During culture, the plate was on the agitator. Spheroids were exposed to 2 mM AGuIX^®^-Cy5.5 nanoparticles at day 3. To avoid losing spheroids, half of the medium was withdrawn and 100 μl of AGuIX^®^-Cy5.5 nanoparticles at a concentration of 4mM were added in each well to have a final concentration of 2mM. After 24 h, spheroids were rinsed with fresh medium (3X, 15 min), fixed with PFA 4%, permeabilized with PBS/0.1% Triton-X, and blocked with PBS/2% BSA/0.1% Triton-X. The spheroids were then labelled with nucgreen™ at a dilution of 1 drop/ 2 ml for overnight at room temperature before being rinsed with PBS (3X, 5 min).

Spheroids could not be imaged in a standard 96-well plate using confocal microscopy; thus they were transferred to an ibidi 96-well plate and imaged.

The same Matlab routine that was used to analyse images of spheroids in microwells was used to analyse these images. Despite the fact that the number of imaged spheroids in this experiment is considerably lower than spheroids in microwells which is due to the limitations of different steps of experiments using a standard 96-well plate, the analysed results show that the distribution and amount of uptaken nanoparticles are similar to spheroids in microwells. This finding supports the permeability of agarose gels for AGuIX^®^-Cy5.5 nanoparticles and validates the usage of such microsystems for spheroids generation and high-throughput drug screening in a more practicable and reproducible manner.

### Figure SI 8. HCT-116 cell incubated in 2D with nanoparticles and spheroid formation afterwards

To make a comparison between cellular uptake of AGuIX^®^-Cy5.5 nanoparticles in monolayer cells and multicellular tumour spheroids, two parallel experiments have been done.

In the first experiment, HCT-116 cells were seeded in agarose-based microwells (**Fig. SI 8A A, Step I**) and after 48 h were exposed to AGuIX^®^-Cy5.5 nanoparticles for 24h (**Fig. SI 8A, Step II**) followed by fixation (**Fig. SI 8A, Step III**).

In the other experiment HCT-116 monolayer cells that were first exposed to AGuIX^®^-Cy5.5 nanoparticles for 24 h (**Fig. SI 8B, Step I**), and then seeded in agarose-based microwells to allow spheroid formation (**Fig. SI 8B, Step II**). After 72h, these spheroids were fixed (**Fig. SI 8B, Step III**).

Spheroids from both experiments were clarified with glycerol 80% (**Fig. SI 8, Step III**) and imaged via confocal microscopy. The confocal images of these two experiments and orthogonal view of spheroids demonstrate that when spheroids are incubated with nanoparticles, clusters could be observed evenly in extracellular and intracellular regions of spheroids and nanoparticles clusters are more in peripheral region than in the centre (**Fig. SI 8A**). When spheroids are made from already AGuIX^®^-Cy5.5 labelled cells, only sparse clusters, with a scattered distribution are observed in spheroids (**Fig. SI 8B)**.

### Figure SI 9. Nanoscale Secondary Ion Mass Spectrometry control analysis

### Figure SI 10. Localization of AGuIX-Cy5.5-nanoparticles after an extensive washing procedure

**Figure SI 1.**
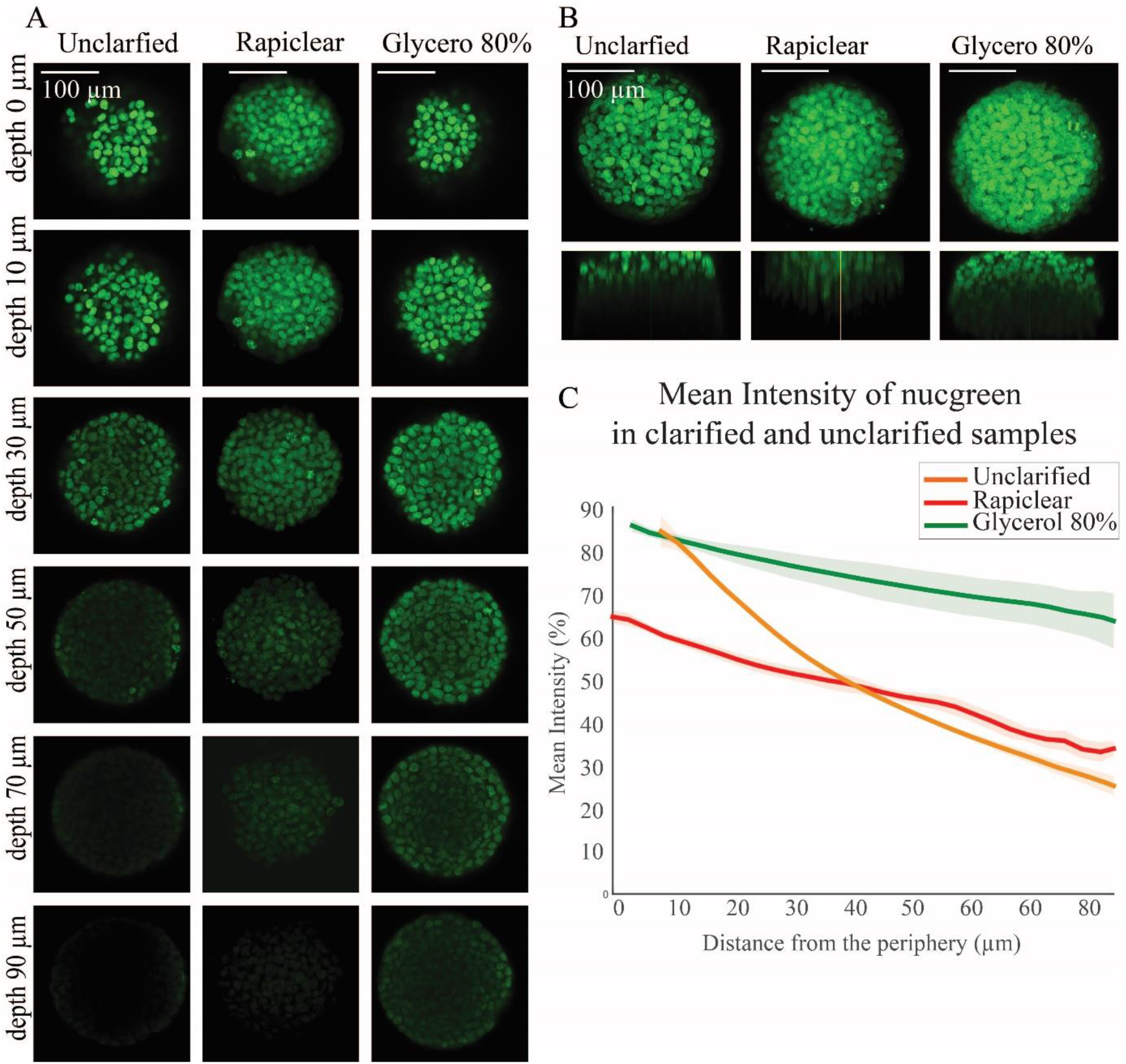
Influence of clarification technique on image acquisition and fluorescence signals. **(A)** Confocal fluorescence images of HCT-116 cell spheroids labelled with Nucgreen in different depth for unclarified, clarified with Rapiclear and clarified with 80%/20% glycerol/PBS respectively. **(B)** Maximal Image Projection (MIP) and xz images of clarified and unclarified spheroids. **(C)** Mean intensity of clarified (green, glycerol-clarification, red, rapiclear-clarification) and unclarified (orange curve) spheroids as a function of the distance from the periphery. Error bars represent standard errors of the Mean (N=10 spheroids for each condition).

**Figure SI 2.**
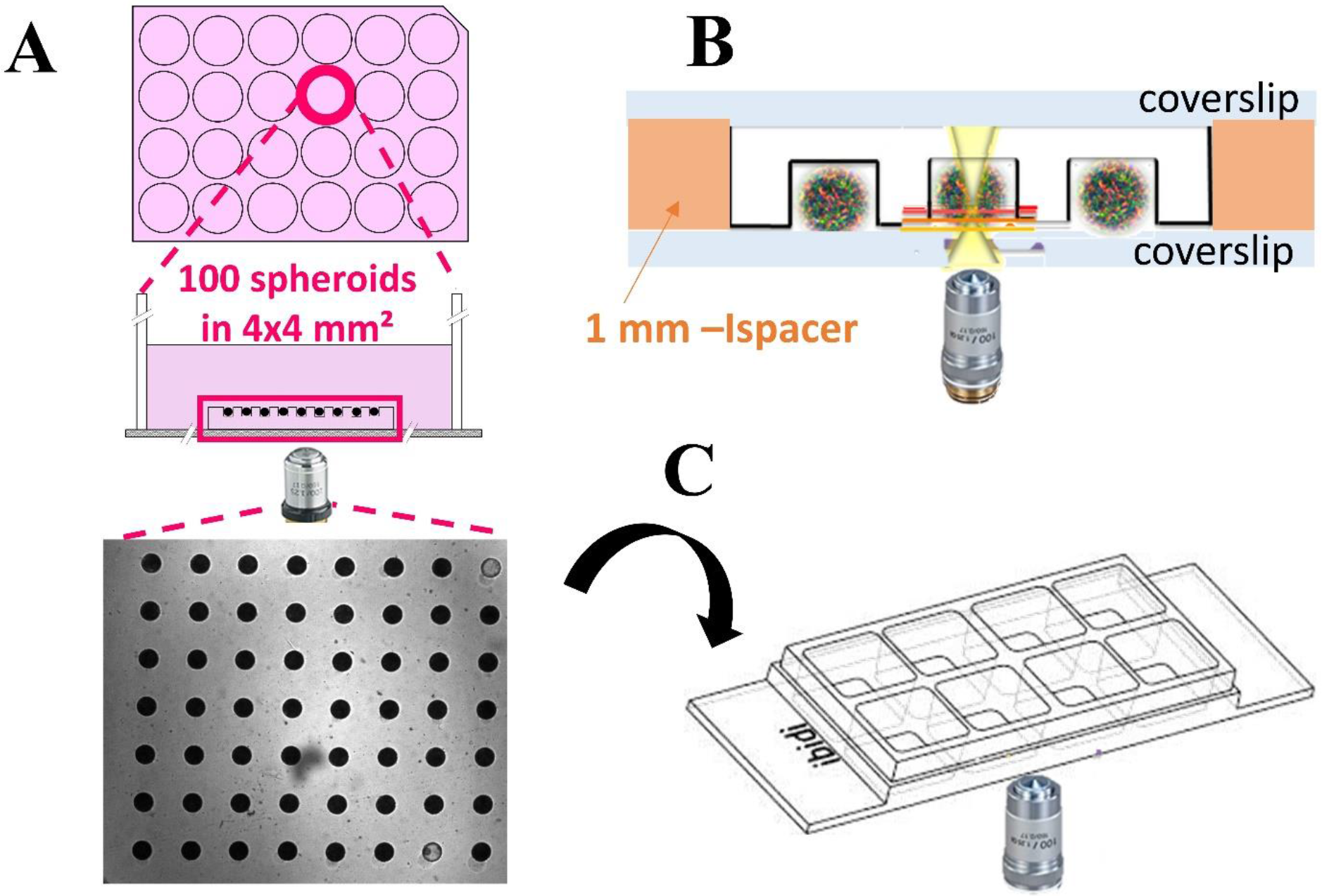
Schematic representation of the mounting procedures used for optical imaging. **(A)** Schematic representation of free-standing microwells, placed on each well of a multi-well plate. **(B)** Schematic representation of the mounting used for optical imaging of fixed samples. The agarose microsystems containing the fixed, immunostained and clarified spheroids are mounted between 2 coverslips using a 1 mm sticky spacer (orange-part, Ispacer from SunjinLab). **(C)** For live imaging of spheroids, it is also possible to transfer the microwells in optical imaging chamber (such as Ibidi^®^ 8-well plates).

**Figure SI 3.**
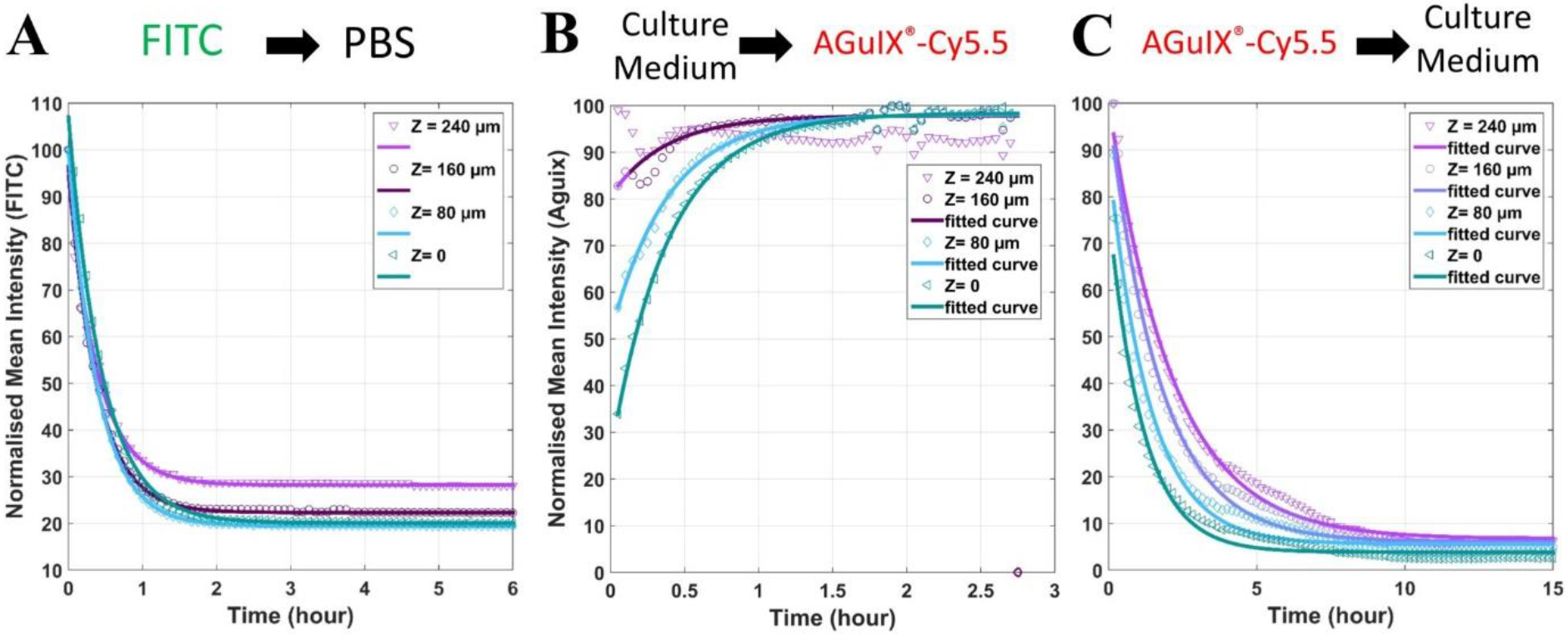
**(A)** Reduction in fluorescence intensity of FITC dye in an agarose microsystem over time for different depth (from Z=0, corresponding to the depth closest to the objective and farthest from the solution reservoir to Z=240 μm, farthest from the objective and closest to solution reservoir). Experimental points are plotted with different markers (Z=0, (pink triangles pointing down), Z=80 μm (blue diamonds), Z=160 μm (purple circles), Z=240 μm (green triangles pointing left)) and the corresponding exponential fit are plotted in bold lines [fitting model a*exp(-time/T)+b].
**(B)** Increase in fluorescence intensity of AGuIX^®^-Cy5.5 nanoparticles in an agarose microsystem over time for different depth (same legend than in (A)). [fitting model a*(1-exp(-time/T))+c].
**(C)** Reduction in fluorescence intensity of AGuIX^®^-Cy5.5 nanoparticles in an agarose microsystem over time for different depth (same legend than in (A). [fitting model a*exp(-time/T)+b].

**Figure SI 4.**
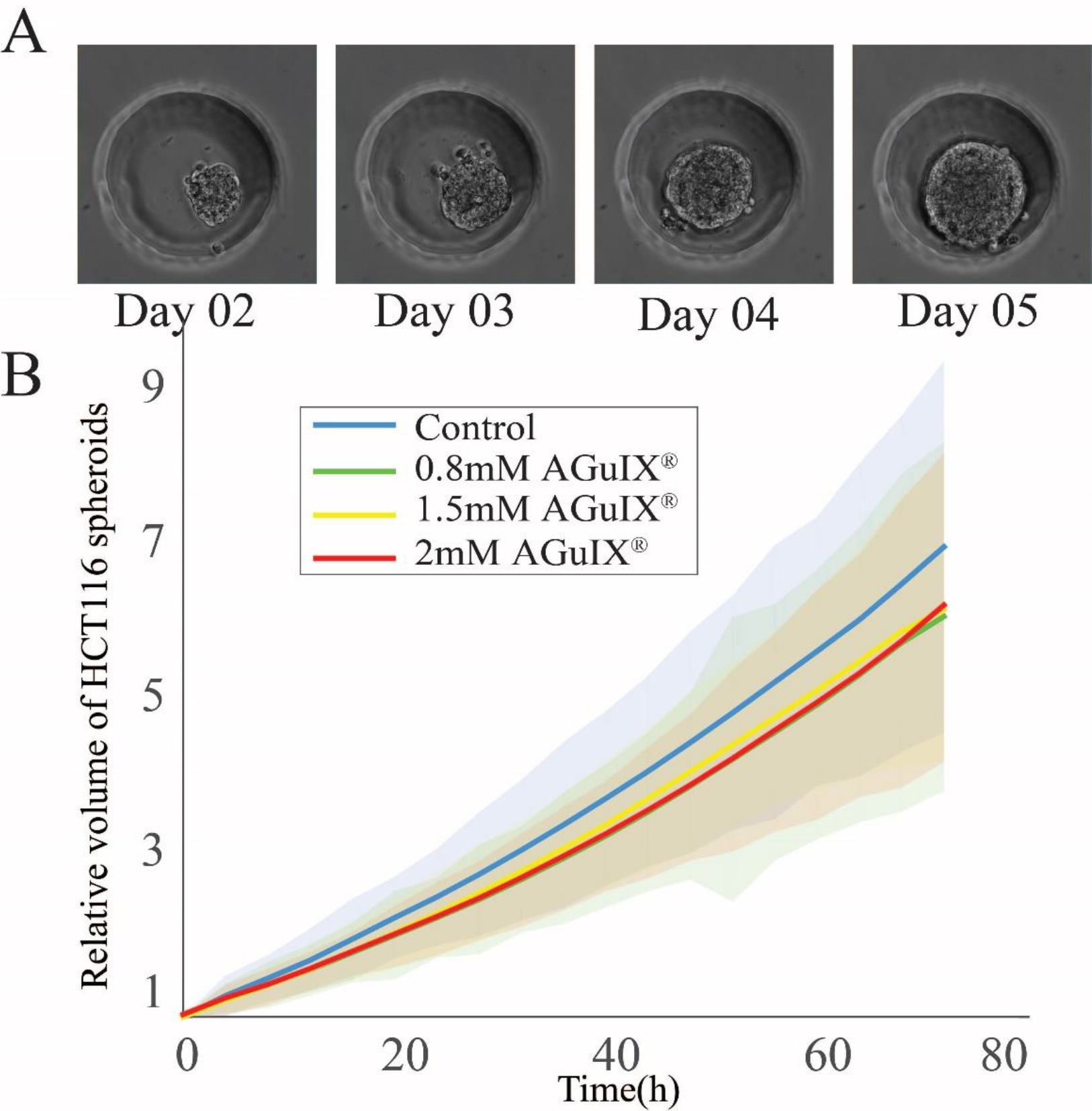
Follow-up of spheroids growth via optical time-lapse microscopy. **(A)** Representative images of daily growth of control HCT-116 spheroids from day 2 to day 5. The well is 200 μm in diameter.
**(B)** Evolution of the relative volume of spheroids as a function of time for control sample and in the presence of three different concentrations of AGuIX^®^-Cy5.5 nanoparticles. Bold lines represent the mean values, and light area represents the standard deviation for each condition (control –blue- [N=102 spheroids], 0.8mM –green-[N=89 spheroids], 1.5mM –yellow-[N=88 spheroids], 2mM –red-[N=102 spheroids]). Three independent experiments for each condition.

**Figure SI 5.**
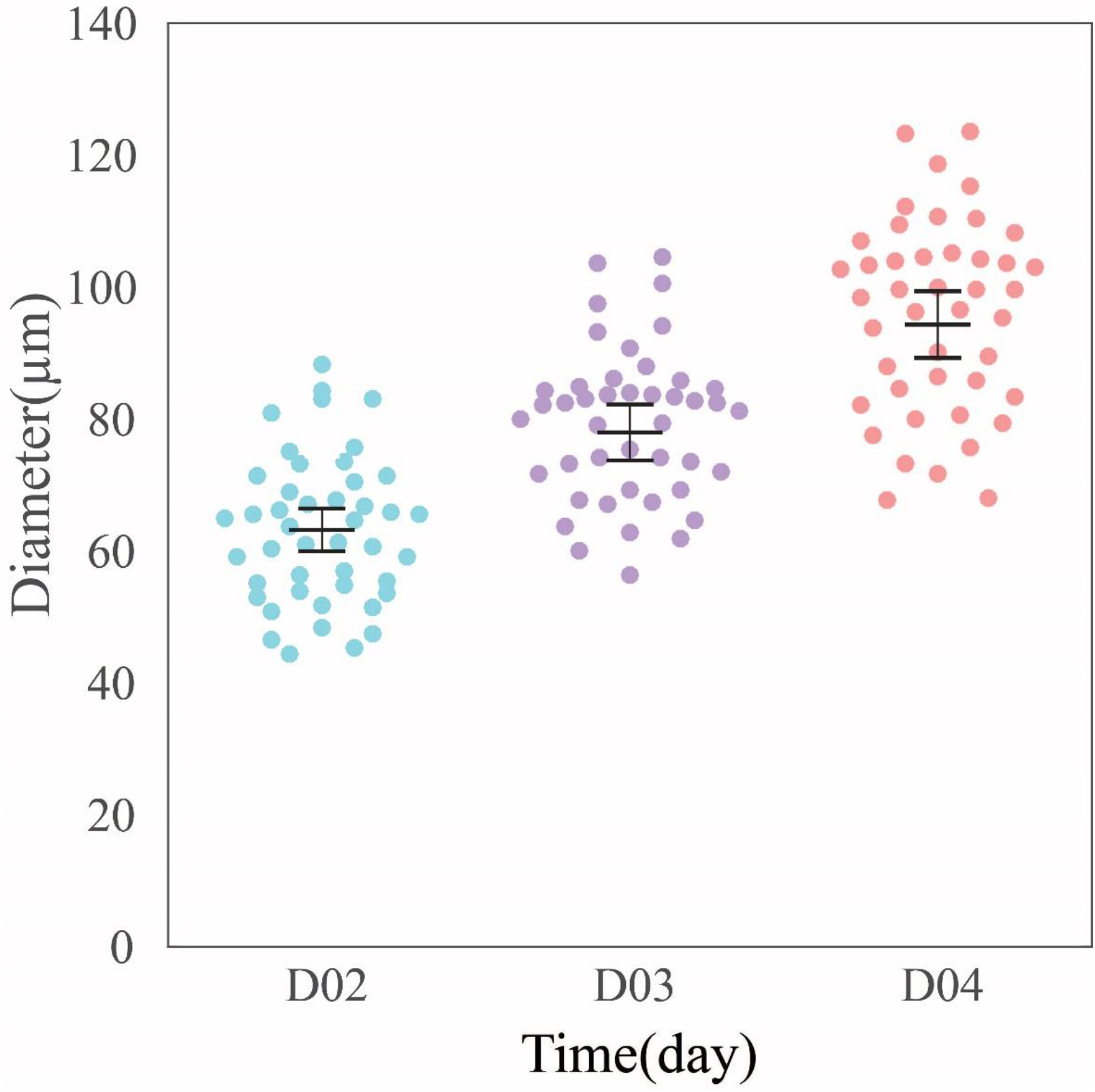
Distribution of HCT-116 multicellular tumour spheroids at day two (blue circles), day three (purple circle) and day four (orange circles) after cell seeding in the agarose-based microwells. Mean values and 95 % Standard Error of the Mean are represented.

**Figure SI 6.**
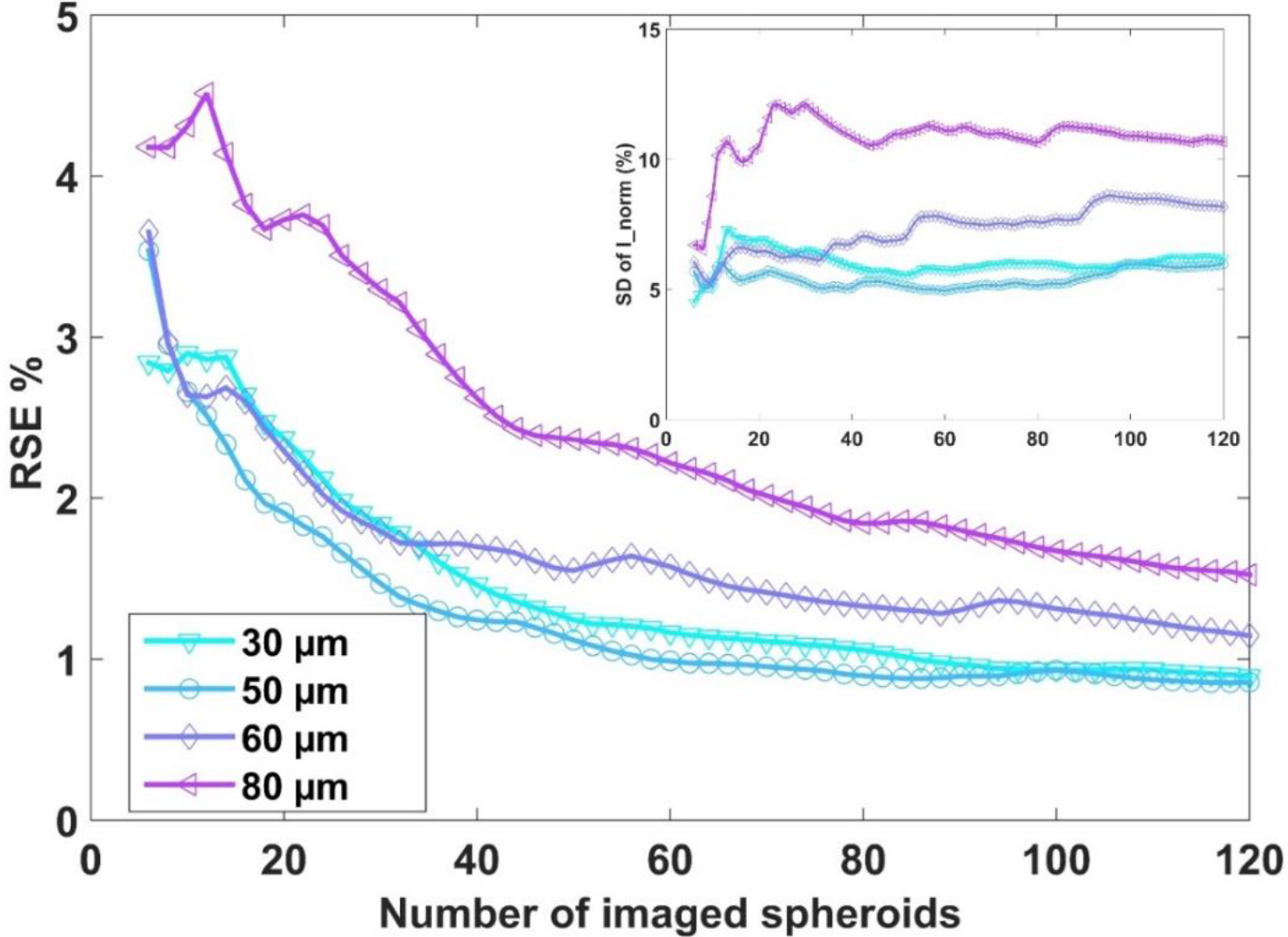
Error analysis due to the number of spheroids analysed. Relative Standard Error (RSE=SEM/mean) as a function of the number of spheroids for different distance from the periphery (30 μm (cyan, triangles pointing down), 50 μm (light blue, circles), 60 μm (intense blue, diamonds) and 80 μm (purple, triangles pointing left). <u>**Inset:**</u> Standard Deviation (SD) of the normalised Intensity I_norm as a function of the number of spheroids, for the same distance from the periphery.

**Figure SI 7.**
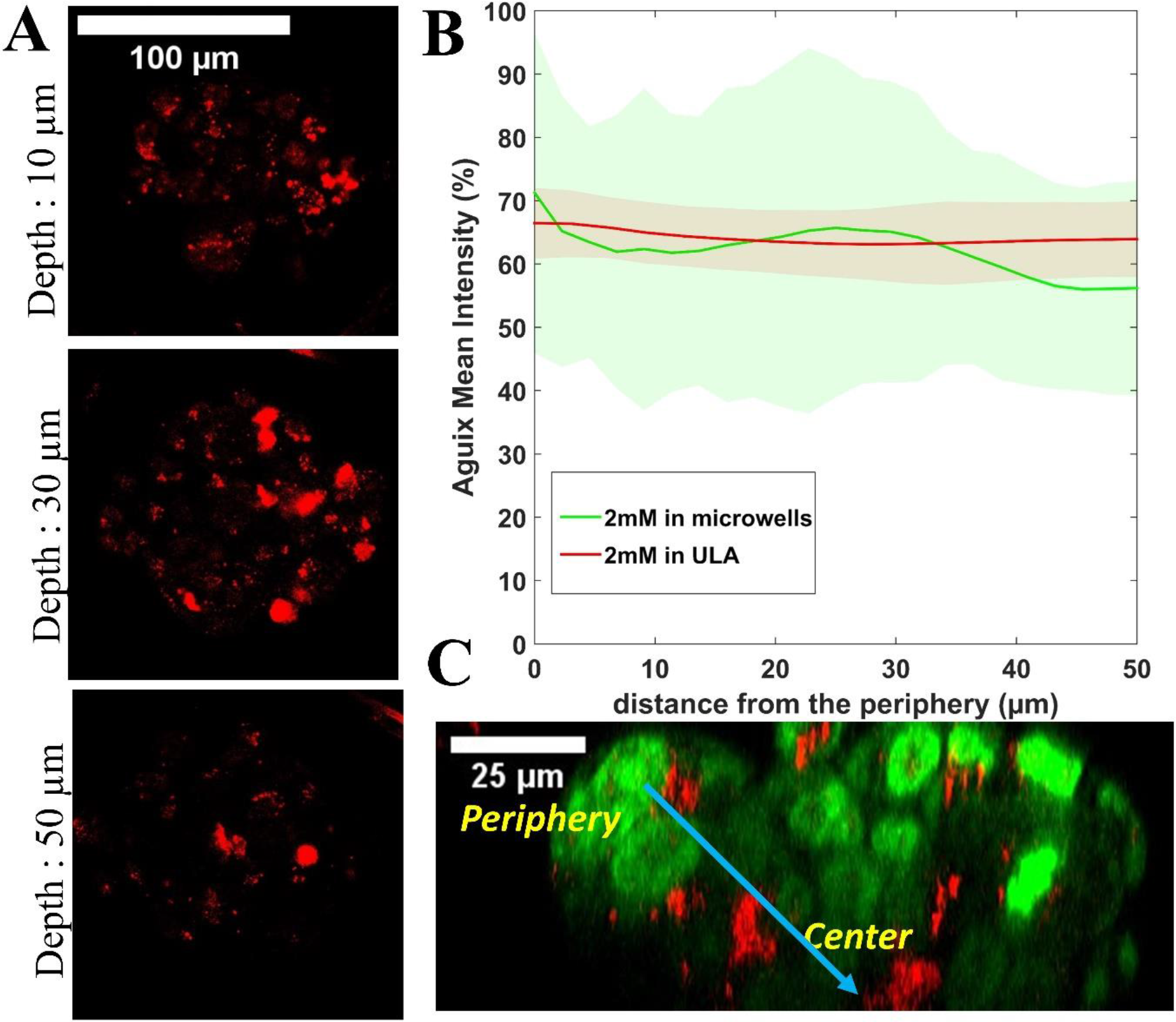
Control experiments using Ultra-Low Adhesion (ULA)multi-well plate. **(A)** Representative confocal fluorescence images of HCT-116 spheroids grown in Ultra-Low-Adhesion multi-well plates for 4 days, then incubated with 2 mM concentration of AGuIX^®^-Cy5.5 for 24 h for three different depths (10, 30 and 50 μm). **(B)** Mean intensity of AGuIX^®^-Cy 5.5 after 24h incubation with 2 mM AGuIX^®^-Cy5.5 as a function of the distance from the periphery in our microsystems (red, N=121, three independent experiment), and in ULA multi-well plates (green, N=10, one experiment). Standard deviations are shown in light colors. **(C)** Orthogonal view of the spheroid in (A) (green=nuclei, red = AGuIX^®^-Cy5.5).

**Figure SI. 8.**
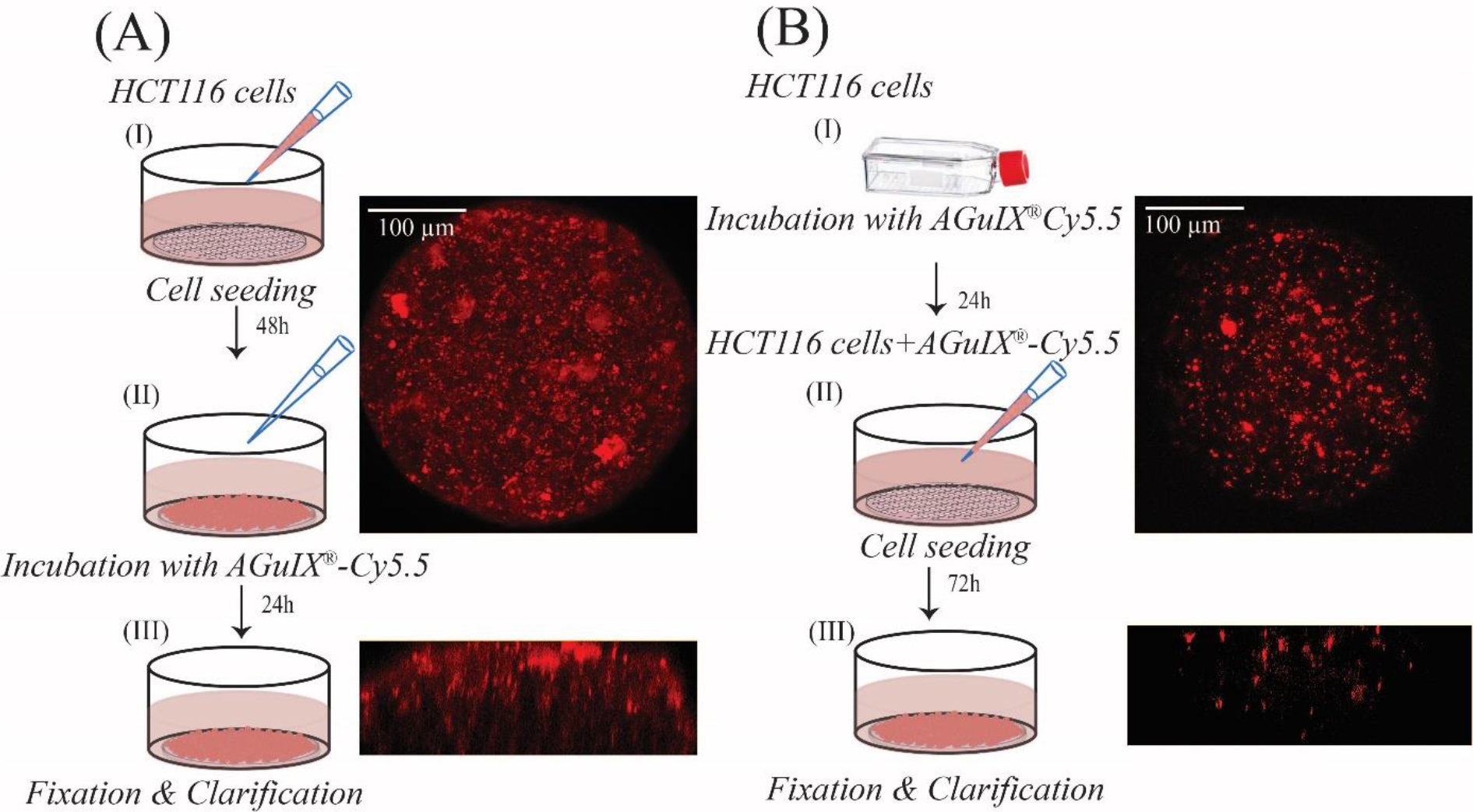
Difference in distribution of AGuIX^®^-Cy 5.5 nanoparticles in HCT116 multicellular tumour spheroids incubated in 2D and 3D cell culture. (A) Spheroids were prepared with agarose-based microwells (**Step I**). After 48 h, they were exposed to 2mM AGuIX^®^-Cy5.5 nanoparticles for 24h **(step II)** and were fixed, clarified **(Step III)** and imaged with confocal microscopy. (B) Monolayer HCT116 cells were first incubated with 2mM AGuIX^®^ nanoparticles for 24h **(Step I)** and then HCT116 spheroids were prepared with these cells **(step II)**. Spheroids were fixed, clarified **(Step III)** and imaged with confocal microscopy.

**Figure SI. 9.**
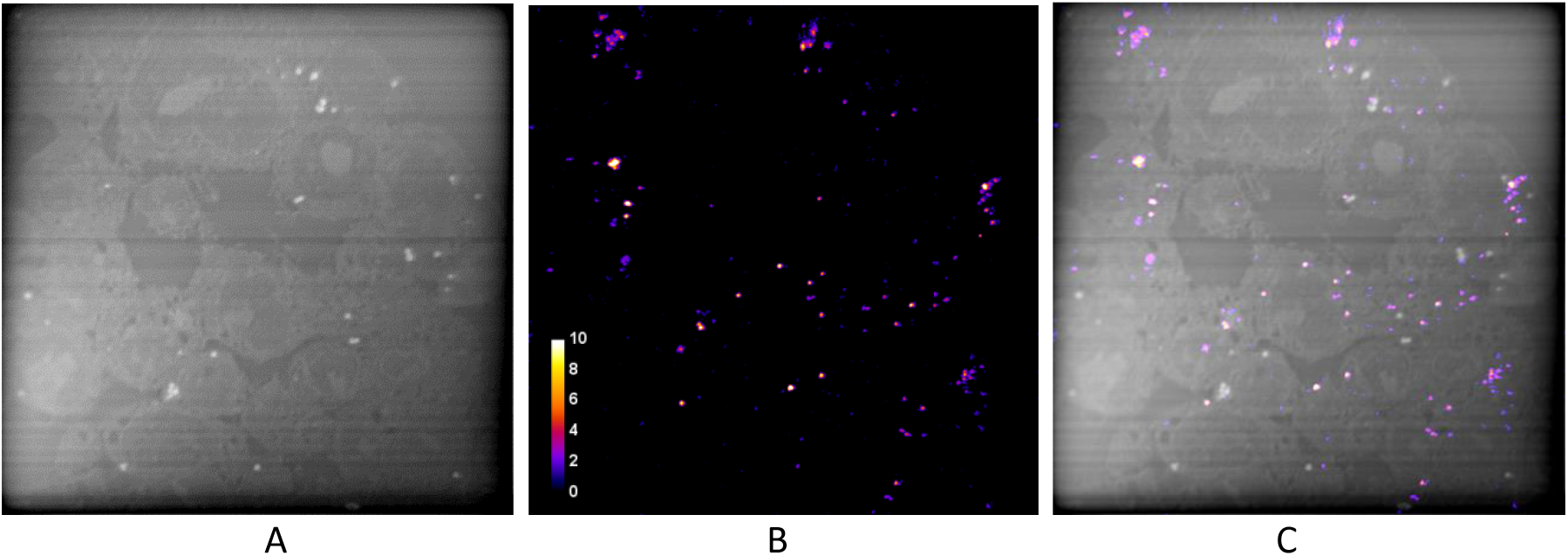
**(A)** NanoSIMS image of ^12^C^−^ of the same area as in **Figure 5** provides the proof of the entirety of the section. The slight contrast is due to the compositional variation of different cells compartments and the surrounding resin (the actual contrast is much lower). A few spots with unusually high ^12^C^−^ emission are probably location of vacuoles. **(B)** Recalled of the distribution of AGuIX^®^-Cy5.5 nanoparticles. **(C)** Merged image of ^28^Si^−^ and ^12^C^−^. Image field: 60 μm.

**Figure SI. 10.**
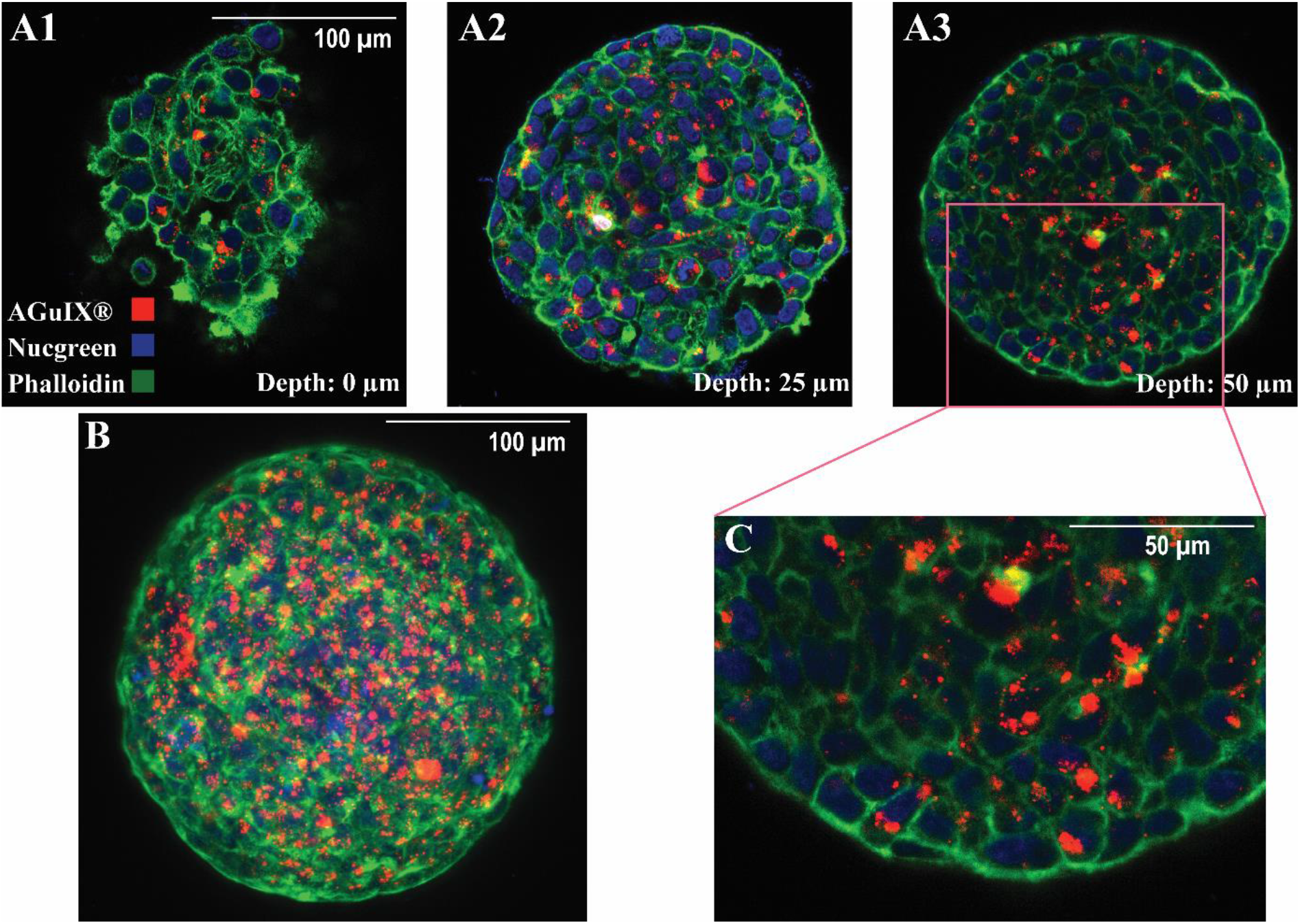
Localization of Aguix-Cy5.5-nanoparticles after an extensive washing procedure within HCT-116 spheroids. Confocal fluorescence images of HCT-116 spheroids incubated with AGuIX^®^-Cy5.5 nanoparticles for 72 h with 2 mM AGuIX^®^-Cy5.5 solution, and washed according to the procedure mentioned in Figure 4, then fixed and immunostained with antibodies to find colocalization of nanoparticles in spheroids. For all images, red, blue and green channels are staining AGuIX^®^-Cy5.5, nuclei and phalloidin (Actins) respectively. **(A1-A3)** Representative images of phalloidin immunostaining merged with AGuIX^®^-Cy5.5 and Nucgreen layers obtained at various depths (**A1-0μm, A2-25 μm, A3-50μm)**. **(B)** Maximal Image Projection (MIP) of confocal fluorescence image of spheroid in (A1-A3). **(C)** Zoomed-in portion of merged image at a depth of 50 μm (square in A3).

## References

1 S. Hua, M. B. C. de Matos, J. M. Metselaar and G. Storm, Front. Pharmacol., 2018, 9, 1–14.

2 L. J. Bray, D. W. Hutmacher and N. Bock, Front. Bioeng. Biotechnol., 2019, 7, 1–36.

3 S. E. Gould, M. R. Junttila and F. J. De Sauvage, Nat. Med., 2015, 21, 431–439.

4 G. Lazzari, P. Couvreur and S. Mura, Polym. Chem., 2017, 8, 4947–4969.

5 A. Sontheimer-Phelps, B. A. Hassell and D. E. Ingber, Nat. Rev. Cancer, 2019, 19, 65–81.

6 S. Peel, A. M. Corrigan, B. Ehrhardt, K. J. Jang, P. Caetano-Pinto, M. Boeckeler, J. E. Rubins, K. Kodella, D. B. Petropolis, J. Ronxhi, G. Kulkarni, A. J. Foster, D. Williams, G. A. Hamilton and L. Ewart, Lab Chip, 2019, 19, 410–421.

7 D. Peer, J. M. Karp, S. Hong, O. C. Farokhzad, R. Margalit and R. Langer, Nat. Nanotechnol., 2007, 2, 751–760.

8 D. Rosenblum, N. Joshi, W. Tao, J. M. Karp and D. Peer, Nat. Commun., 2018, 9, 1410.

9 M. Zanoni, F. Piccinini, C. Arienti, A. Zamagni, S. Santi, R. Polico, A. Bevilacqua and A. Tesei, Sci. Rep., 2016, 6, 1–11.

10 S. Wilhelm, A. J. Tavares, Q. Dai, S. Ohta, J. Audet, H. F. Dvorak and W. C. W. Chan, Nat. Rev. Mater., 2016, 1, 16014.

11 E. J. Guggenheim, S. Milani, P. J. F. Röttgermann, M. Dusinska, C. Saout, A. Salvati, J. O. Rädler and I. Lynch, NanoImpact, 2018, 10, 121–142.

12 W. Asghar, R. El Assal, H. Shafiee, S. Pitteri, R. Paulmurugan and U. Demirci, Mater. Today, 2015, 18, 539–553.

13 S. Nath and G. R. Devi, Pharmacol. Ther., 2016, 163, 94–108.

14 M. Millard, I. Yakavets, V. Zorin, A. Kulmukhamedova, S. Marchal and L. Bezdetnaya, Int. J. Nanomedicine, 2017, 12, 7993–8007.

15 F. Hirschhaeuser, H. Menne, C. Dittfeld, J. West, W. Mueller-klieser and L. A. Kunz-schughart, J. Biotechnol., 2010, 148, 3–15.

16 H. L. Ma, Q. Jiang, S. Han, Y. Wu, J. C. Tomshine, D. Wang, Y. Gan, G. Zou and X. J. Liang, Mol. Imaging, 2012, 11, 487–498.

17 S. Huo, H. Ma, K. Huang, J. Liu, T. Wei, S. Jin, J. Zhang, S. He and X. J. Liang, Cancer Res., 2013, 73, 319–330.

18 A. Virgone-Carlotta, M. Lemasson, H. C. Mertani, J. J. Diaz, S. Monnier, T. Dehoux, H. Delanoë-Ayari, C. Rivière and J. P. Rieu, PLoS One, 12(11):e0188100.

19 B. Rodday, F. Hirschhaeuser, S. Walenta and W. Mueller-Klieser, J. Biomol. Screen., 2011, 16, 1119–1124.

20 J. M. Kelm, N. E. Timmins, C. J. Brown, M. Fussenegger and L. K. Nielsen, Biotechnol. Bioeng., 2003, 83, 173–180.

21 Y.-C. Chen, P. N. Ingram, S. Fouladdel, S. P. McDermott, E. Azizi, M. S. Wicha and E. Yoon, Sci. Rep., 2016, 6, 27301.

22 M. Akay, J. Hite, N. G. Avci, Y. Fan, Y. Akay, G. Lu and J. J. Zhu, Sci. Rep., 2018, 8, 1–9.

23 R. Mukhopadhyay, Anal. Chem., 2007, 79, 3249–3253.

24 B. J. van Meer, H. de Vries, K. S. A. Firth, J. van Weerd, L. G. J. Tertoolen, H. B. J. Karperien, P. Jonkheijm, C. Denning, A. P. IJzerman and C. L. Mummery, Biochem. Biophys. Res. Commun., 2017, 482, 323–328.

25 M. W. Toepke and D. J. Beebe, Lab Chip, 2006, 6, 1484–1486.

26 D. T. Butcher, T. Alliston and V. M. Weaver, Nat. Rev. Cancer, 2009, 9, 108–22.

27 J. M. Lee, D. Y. Park, L. Yang, E. J. Kim, C. D. Ahrberg, K. B. Lee and B. G. Chung, Sci. Rep., 2018, 8, 1–10.

28 Y. Li and E. Kumacheva, Sci. Adv., 2018, 4, 1–11.

29 X. Gong, C. Lin, J. Cheng, J. Su, H. Zhao, T. Liu, X. Wen and P. Zhao, PLoS One, 2015, 10, e0130348.

30 J. Dahlmann, G. Kensah, H. Kempf, D. Skvorc, A. Gawol, D. A. Elliott, G. Dräger, R. Zweigerdt, U. Martin and I. Gruh, Biomaterials, 2013, 34, 2463–2471.

31 D. L. Priwitaningrum, J. B. G. Blondé, A. Sridhar, J. van Baarlen, W. E. Hennink, G. Storm, S. Le Gac and J. Prakash, J. Control. Release, 2016, 244, 257–268.

32 G. Fang, H. Lu, A. Law, D. Gallego-Ortega, D. Jin and G. Lin, Lab Chip, 2019, 19, 4093–4103.

33 X. Hu, X. Hu, S. Zhao, S. Zhao, Z. Luo, Y. Zuo, Y. Zuo, F. Wang, F. Wang, J. Zhu, J. Zhu, L. Chen, L. Chen, D. Yang, Y. Zheng, Y. Zheng, Y. Cheng, F. Zhou, Y. Yang and Y. Yang, Lab Chip, 2020, 20, 2228–2236.

34 V. Normand, D. L. Lootens, E. Amici, K. P. Plucknett and P. Aymard, Biomacromolecules, 2000, 1, 730–738.

35 T. H. Jovic, G. Kungwengwe, A. C. Mills and I. S. Whitaker, Front. Mech. Eng., 2019, 5, 19.

36 A. Pluen, P. A. Netti, R. K. Jain and D. A. Berk, Biophys. J., 1999, 77, 542–52.

37 F. Lux, V. L. Tran, E. Thomas, S. Dufort, F. Rossetti, M. Martini, C. Truillet, T. Doussineau, G. Bort, F. Denat, F. Boschetti, G. Angelovski, A. Detappe, Y. Crémillieux, N. Mignet, B. T. Doan, B. Larrat, S. Meriaux, E. Barbier, S. Roux, P. Fries, A. Müller, M. C. Abadjian, C. Anderson, E. Canet-Soulas, P. Bouziotis, M. Barberi-Heyob, C. Frochot, C. Verry, J. Balosso, M. Evans, J. Sidi-Boumedine, M. Janier, K. Butterworth, S. Mcmahon, K. Prise, M. T. Aloy, D. Ardail, C. Rodriguez-Lafrasse, E. Porcel, S. Lacombe, R. Berbeco, A. Allouch, J. L. Perfettini, C. Chargari, E. Deutsch, G. Le Duc and O. Tillement, Br. J. Radiol., 2019, 92, 109320180365.

38 W. Rima, L. Sancey, M. T. Aloy, E. Armandy, G. B. Alcantara, T. Epicier, A. Malchère, L. Joly-Pottuz, P. Mowat, F. Lux, O. Tillement, B. Burdin, A. Rivoire, C. Boulé, I. Anselme-Bertrand, J. Pourchez, M. Cottier, S. Roux, C. Rodriguez-Lafrasse and P. Perriat, Biomaterials, 2013, 34, 181–195.

39 C. Riviere, A. Prunet, L. Fuoco, H. Delanoë-Ayari, Patent FR3079524A1, 2018

40 G. Le Duc, S. Roux, A. Paruta-Tuarez, S. Dufort, E. Brauer, A. Marais, C. Truillet, L. Sancey, P. Perriat, F. Lux and O. Tillement, Cancer Nanotechnol., 2014, 5, 1–14.

41 C. A. Schneider, W. S. Rasband and K. W. Eliceiri, Nat. Methods, 2012, 9, 671–675.

42 L. Le Roux, A. Volgin, D. Maxwell, K. Ishihara, J. Gelovani and D. Schellingerhout, Mol. Imaging, 2008, 7, 214–221.

43 J. F. Dekkers, M. Alieva, L. M. Wellens, H. C. R. Ariese, P. R. Jamieson, A. M. Vonk, G. D. Amatngalim, H. Hu, K. C. Oost, H. J. G. Snippert, J. M. Beekman, E. J. Wehrens, J. E. Visvader, H. Clevers and A. C. Rios, Nat. Protoc., 2019, 14, 1756–1771.

44 T. Silva Santisteban, O. Rabajania, I. Kalinina, S. Robinson and M. Meier, Lab Chip, 2018, 18, 153–161.

45 E. Nürnberg, M. Vitacolonna, J. Klicks, E. von Molitor, T. Cesetti, F. Keller, R. Bruch, T. Ertongur-Fauth, K. Riedel, P. Scholz, T. Lau, R. Schneider, J. Meier, M. Hafner and R. Rudolf, Front. Mol. Biosci., 2020, 7, 1–19.

46 J. L. Guerquin-Kern, T. Di Wu, C. Quintana and A. Croisy, Biochim. Biophys. Acta - Gen. Subj., 2005, 1724, 228–238.

47 G. Slodzian, B. Daigne, F. Girard, F. Boust and F. Hillion, Biol. Cell, 1992, 74, 43–50.

48 C. MessaoudiI, T. Boudier, C. O. S. Sorzano and S. Marco, BMC Bioinformatics, 2007, 8, 1–9.

49 M. Singh, D. A. Close, S. Mukundan, P. A. Johnston and S. Sant, Assay Drug Dev. Technol., 2015, 13, 570–583.

50 L. B. Sims, L. T. Curtis, H. B. Frieboes and J. M. Steinbach-Rankins, J. Nanobiotechnology, 2016, 14, 1–12.

51 A. R. Kang, H. I. Seo, B. G. Chung and S. H. Lee, Nanomedicine Nanotechnology, Biol. Med., 2015, 11, 1153–1161.

52 L. Štefančíková, E. Porcel, P. Eustache, S. Li, D. Salado, S. Marco, J. L. Guerquin-Kern, M. Réfrégiers, O. Tillement, F. Lux and S. Lacombe, Cancer Nanotechnol., 2014, 5, 1–15.

53 L. Sancey, F. Lux, S. Kotb, S. Roux, S. Dufort, A. Bianchi, Y. Crémillieux, P. Fries, J.-L. Coll, C. Rodriguez-Lafrasse, M. Janier, M. Dutreix, M. Barberi-Heyob, F. Boschetti, F. Denat, C. Louis, E. Porcel, S. Lacombe, G. Le Duc, E. Deutsch, J.-L. Perfettini, A. Detappe, C. Verry, R. Berbeco, K. T. Butterworth, S. J. McMahon, K. M. Prise, P. Perriat and O. Tillement, Br. J. Radiol., 2014, 87, 20140134.

54 T. Stylianopoulos, L. L. Munn and R. K. Jain, Trends in Cancer, 2018, 4, 292–319.

55 K. Carver, X. Ming and R. L. Juliano, Mol. Ther. - Nucleic Acids, 2014, 3, e153.

56 H. Kang, S. Mintri, A. V. Menon, H. Y. Lee, H. S. Choi and J. Kim, Nanoscale, 2015, 7, 18848–18862.

57 K. Raza, P. Kumar, N. Kumar and R. Malik, Pharmacokinetics and biodistribution of the nanoparticles, Elsevier Ltd, 2017.

58 V. Ivošev, G. J. Sánchez, L. Stefancikova, D. A. Haidar, C. R. González Vargas, X. Yang, R. Bazzi, E. Porcel, S. Roux and S. Lacombe, Nanotechnology, 2020, 31, 13.

59 N. D. Donahue, H. Acar and S. Wilhelm, Adv. Drug Deliv. Rev., 2019, 143, 68–96.

60 C. A. Lyssiotis and A. C. Kimmelman, Trends Cell Biol., 2017, 27, 863–875.

61 S. Behzadi, V. Serpooshan, W. Tao, M. A. Hamaly, M. Y. Alkawareek, E. C. Dreaden, D. Brown, A. M. Alkilany, O. C. Farokhzad and M. Mahmoudi, Chem. Soc. Rev., 2017, 46, 4218–4244.

62 J. Kondo, T. Ekawa, H. Endo, K. Yamazaki, N. Tanaka, Y. Kukita, H. Okuyama, J. Okami, F. Imamura, M. Ohue, K. Kato, T. Nomura, A. Kohara, S. Mori, S. Dan and M. Inoue, Cancer Sci., 2019, 110, 345–355.

63 S. E. Park, A. Georgescu and D. Huh, Science (80-.)., 2019, 364, 960–965.

64 A. Ahmed, S. Goodarzi, C. Frindel, G. Recher, C. Riviere and D. Rousseau, bioRxiv, , DOI:https://doi.org/10.1101/2021.01.31.428996.

65 A. Prunet, S. Lefort, H. Delanoë-Ayari, B. Laperrousaz, G. Simon, C. Barentin, S. Saci, F. Argoul, B. Guyot, J.-P. Rieu, S. Gobert, V. Maguer-Satta and C. Rivière, Lab Chip, 2020, 20, 4016–4030.

66 I. F. Rizzuti, P. Mascheroni, S. Arcucci, Z. Ben-Mériem, A. Prunet, C. Barentin, C. Rivière, H. Delanoë-Ayari, H. Hatzikirou, J. Guillermet-Guibert and M. Delarue, Phys. Rev. Lett., 2020, 125, 128103.

67 A. Albanese, A. K. Lam, E. A. Sykes, J. V. Rocheleau and W. C. W. Chan, Nat. Commun., 2013, 4, 1–8.

68 T. Yu, J. Zhu, Y. Li, Y. Ma, J. Wang, X. Cheng, S. Jin, Q. Sun, X. Li, H. Gong, Q. Luo, F. Xu, S. Zhao and D. Zhu, Sci. Rep., 2018, 8, 1–9.

